# Credible set is sensitive to imputation quality and missing variants

**DOI:** 10.1101/2024.08.28.610135

**Authors:** Yanyu Liang, 23andMe Research Team, Adam Auton, Xin Wang

## Abstract

Bayesian fine-mapping to obtain credible sets has been widely applied post GWAS to pinpoint causal variants. The calculation of credible sets generally assumes that all variants have been equally well genotyped, which is often not the case when a GWAS has been run on imputed data. In this work, we investigate the behavior of credible sets in imputed datasets utilizing ‘held out’ genotyped variants to measure accuracy. We show, via simulation, that: i) the coverage of credible sets decreases when using imputed variants in GWAS; ii) rare causal variants often fail to be tagged in credible sets when they are not present in the GWAS variant set. We develop a reweighting approach to take imputation quality into account during fine-mapping that only requires summary statistics, and demonstrate the approach with real data.

## 1 Introduction

Statistical fine-mapping has been widely adopted for identifying causal variants in genome-wide association studies (GWAS). Many fine-mapping methods (Wellcome Trust Case Control Consortium et al. 2012; Hormozdiari et al. 2014; Benner et al. 2016; Wen et al. 2016; Wang et al. 2020) rely on a Bayesian variable selection framework, in which a prior probability of being causal is assigned to each variant and the method returns the corresponding posterior probability of each variant being causal (i.e. posterior inclusion probability, henceforth PIP) after observing GWAS data. On the basis of PIPs, fine-mapping analysis also returns an *x*% credible set which is the minimal set of variants that contain the causal variant with at least *x*% posterior probability.

Existing fine-mapping methods usually make the simplifying assumption that genotypes are observed without error. However, most current GWAS are performed using imputed genotype dosages. This discrepancy introduces challenges in interpreting fine-mapping results. In this work, we consider a common situation in which SNP array genotype calls are available for a subset of imputed dosages, and utilize SNP array genotype calls that were not included within the imputation procedure to assess the performance of credible sets. We show, via simulation, that the quality of Bayesian fine-mapping is affected by imputation quality and credible sets can be not well calibrated, especially when the causal variant is common and imputation quality is poor. To address this problem, we propose a solution that explicitly takes imputation quality into account that only requires summary statistics, and showcase the solution in real GWAS data.

## 2 Results

### Overview of simulation study

To evaluate the influence of genotype uncertainties in fine-mapping, we designed a simulation study (Figure 1; Methods). Leveraging 23andMe, Inc SNP array genotypes and imputed genotype dosages, the simulation used a set of 402 variants that have both genotyped and imputed values. Importantly, these genotyped variants were not included within the input to the imputation methodology, so they are a ‘held out’ set that can be used to measure imputation accuracy. This set of variants represented minor allele frequencies (MAFs) from 1e-6 to 0.49, and despite on average poorer imputation quality for rarer variants, we sampled a range of imputation qualities for rare and common variants (Figure S1). The intent of the variant selection is to include similar numbers of variants within each MAF and imputation quality bin so that the results are not dominated by a particular MAF and/or imputation quality regime. As a result, this variant set is not representative of the genome-wide distribution of imputation quality. For each variant (i.e. the focal variant), our simulation procedure is outlined below (see more details in Methods). We conducted the simulation in 2 million samples of European genetic ancestry (Durand et al. 2021). We simulated a phenotype *y* from *y* = *X* + ϵ, ϵ *∼ N*(0, σ^2^) where *X* is the observed genotype (obtained from a genotyping array) of the focal variant. For the purpose of this simulation, the effect size is set to 1 without loss of generality. To capture a wide range of GWAS power, the value of σ^2^ is set so that 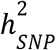 heritability values ranging from 5e-6 to 1e-4 are used, which correspond to expected p-values ranging from 2e-3 to 2e-45. We next performed association testing and fine-mapping analysis (Wellcome Trust Case Control Consortium et al. 2012) within a ±100kb window around the focal variant assuming a single causal variant (the focal variant) where the genotype of the focal variant was: i) microarray genotyped values; ii) imputed dosages; iii) missing. For each of the focal variants, the simulation was repeated 5 times for each 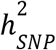 value.

**Figure 1:**
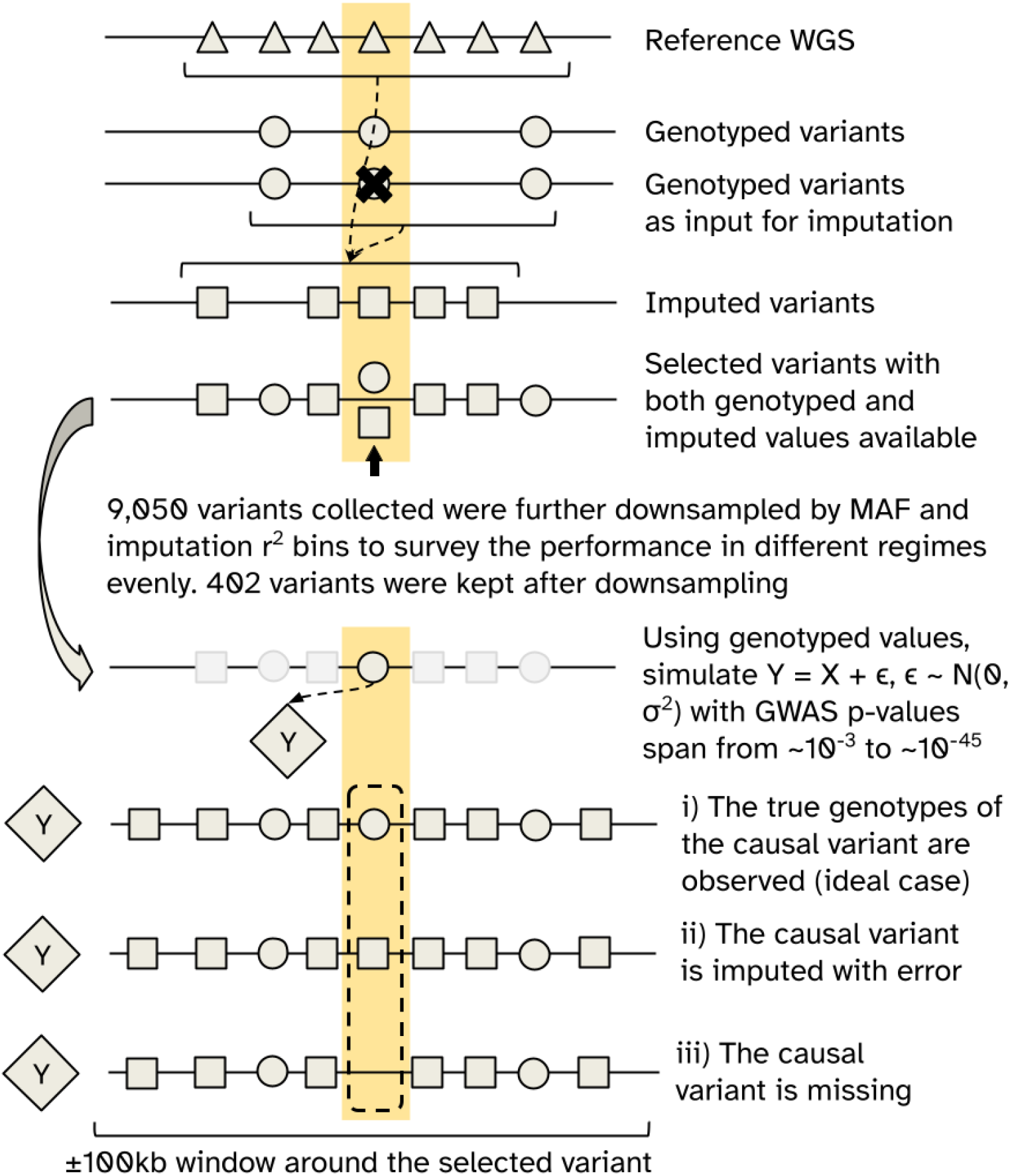
Overview of the simulation study. Variants with both genotyped and imputed dosages were used as the focal variants in the simulation (highlighted in yellow). The genotype values of the focal variants were not included in the imputation procedure, allowing inference of imputation performance.

### Imputation attenuates GWAS marginal statistics

To first examine the effect of genotype uncertainties on the marginal statistics, we compared the association results of the focal variant when using genotyped versus imputed values. As has been shown before (Spencer et al. 2009), the following relationship holds comparing the expected *χ*^2^ statistic when imputed dosages or genotypes are used:

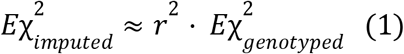

with *r*^2^ being the imputation accuracy (defined as the squared correlation of imputed and true genotypes) (Supplementary Notes 2.1). In our simulation results, we observed concordance between 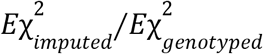 and the imputation quality (Figure S2).

Consistent with this attenuation bias when imputed dosages were used, the estimated effect sizes were attenuated towards zero especially for poorly imputed variants (Figure S3), and the p-value significance is reduced with the amount of reduction correlating with imputation quality (Figure S4). Note that the attenuated *Eχ*^2^ due to imputation resembles the effect of linkage disequilibrium (LD) (Zhu and Stephens 2017):

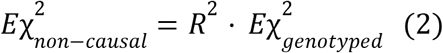

where *R*^2^ captures LD (measured via squared correlation) between the causal variant and a nearby non-causal variant.

### Imputation leads to miscalibrated credible sets

We evaluated the quality of credible sets via: i) the observed coverage of the credible set, ii) calibration of the posterior inclusion probabilities for the focal variant. We first looked into the observed coverage of the 95% credible set, i.e. the fraction of experiments in which the 95% credible set contains the causal variant (Figure 2). When the focal variant is genotyped, the 95% credible set captured the causal variant in *≥* 95% of the experiments. In contrast, when the focal variant is imputed, the observed coverage was *≥* 95% when power is low (i.e.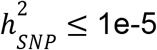, median p-value *≥* 2.1e-17) but gradually decreased, as the association power gets stronger, to 89.7% at the highest heritability/power simulated (i.e.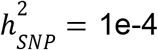, median p-value = 1.6e-33). The observed coverage was, on average, lower when the causal variant is common (MAF > 0.01) than when they are more rare (MAF < 0.01) and the difference in the observed coverage became more apparent if we contrasted causal variants by their maximum LD within the locus (Figure S5). The results were consistent for the observed PIPs of the focal variant (Figures S6 and S7). The PIPs were anti-conservative when imputation uncertainties were present, and the observed fractions of causal variants were significantly below the reported PIPs for strong associations (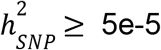, median p-value *≤* 2.1e-17).

**Figure 2:**
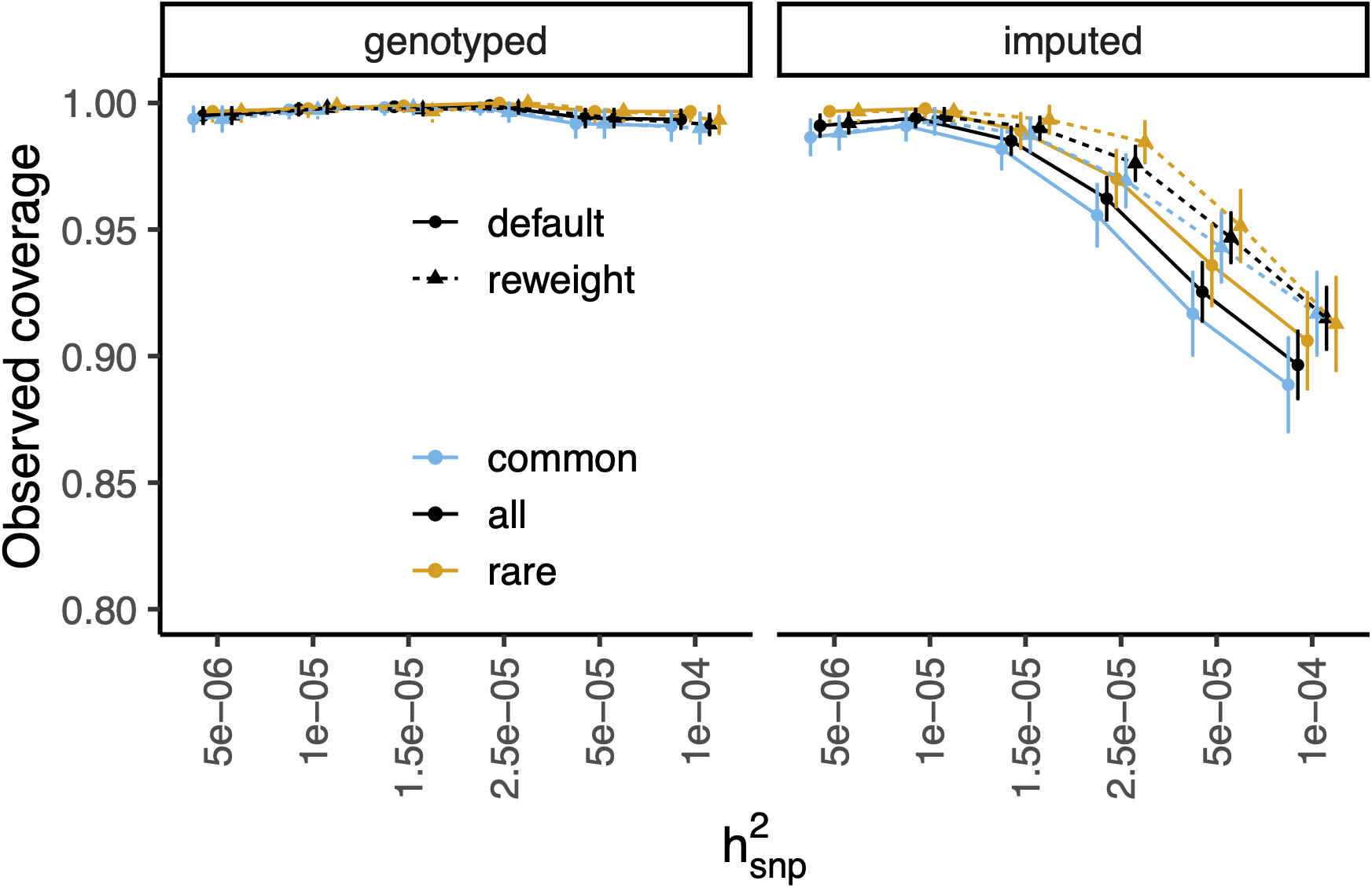
The observed coverage of 95% credible sets when the focal variant is genotyped or imputed. At each 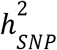 (x-axis), the observed credible set coverage (y-axis) stratified by MAF bins (common: MAF > 0.01; rare: MAF < 0.01) is shown. The two panels show results based on genotyped and imputed values for the focal variant. In each panel, solid lines are results based on the association result as-is. Dashed lines show results from the proposed “reweighting” approach.

### Credible sets fail to tag the causal variant when the causal variant is missing

A critical assumption in Bayesian fine-mapping methods is that the causal variant is typed and included in the model. Next, we examined the scenario where this assumption is violated, i.e. the causal variant is missing in the fine-mapping analysis. We quantified the performance of fine-mapping via the “tagging ability” of the 95% credible set, defined as the median LD (squared genotype correlation) between the causal variant and all variants in the credible set. For comparison, we also calculated this quantity for scenarios where the causal variant was either genotyped or imputed, in which the tagging ability is 1 if the 95% credible set contained only the causal variant. As expected, the tagging ability of the credible set increased as the focal SNP heritability increased (Figure 3 and Figure S8). For a locus at typical GWAS significance (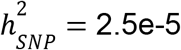, median p-value = 4.6e-9), when a common causal variant (MAF > 0.01) is missing from fine-mapping, the average tagging ability of a 95% credible set decreased from median LD *r*^2^= 0. 69 when the causal variant is imputed, to median LD *r*^2^ = 0. 41. For rare variants (MAF < 0.01), even the most well-powered simulated loci (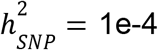, median p-value = 1.6e-33) saw the tagging ability decrease from median *r*^2^ = 0. 86 to median *r*^2^ = 0. 40. On the other hand, when the causal variant is included as an imputed variant, the diminished tagging ability of the 95% credible sets due to imputation can be largely overcome with increased association power, which in practice can be achieved via improved imputation and/or increased sample size. Although imputation uncertainties affect the ability of the credible set to capture a causal variant, the inclusion of the imputed variants can substantially improve the ability of the credible set to tag a causal variant. The improvement is most prominent when the causal variant is rare which suggests substantial value of imputing rare variants in post-GWAS fine-mapping analyses.

**Figure 3:**
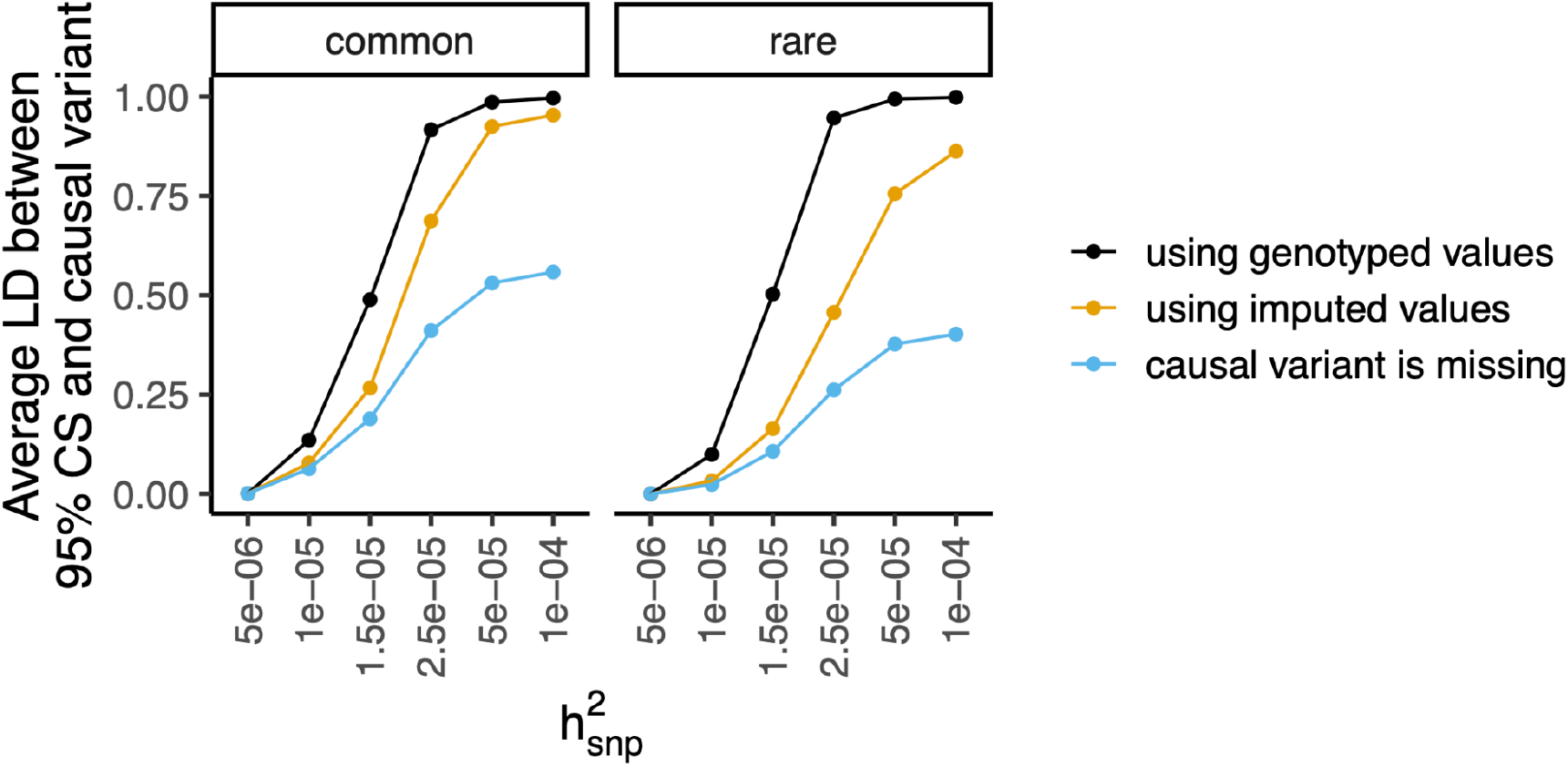
The average LD between the 95% credible set (CS) and the causal variant. For each experiment, fine-mapping analysis is run within the locus with: i) genotyped values at the focal variant (black); ii) imputed values at the focal variant (yellow); iii) the focal variant being missing (blue). To measure the quality of the 95% credible set, the median of LD (the squared correlation) between all the variants among the 95% credible set and the focal variant is calculated. At each 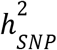 (x-axis), the average value of the median LD across all experiments (y-axis) are shown for common focal variants (MAF > 0.01) and rare focal variants (MAF < 0.01) separately.

### Reweighted fine-mapping improves credible set coverage

To explore methods that account for imputation error in the fine-mapping procedure, we considered the following model:

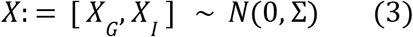

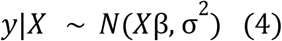

where *X* represents true genotypes. 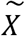 represents observed dosages. Subscripts *G* and *I* indicate variants being genotyped and imputed so that 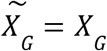. We assume 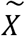 and *X* are standardized and σ is the LD matrix of the locus. Note that Equation 4 is commonly assumed in fine-mapping methods (Wellcome Trust Case Control Consortium et al. 2012; Hormozdiari et al. 2014; Benner et al. 2016; Wen et al. 2016; Wang et al. 2020).

To model imputation error explicitly, we introduce

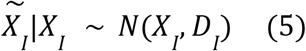

Where *D* is a diagonal matrix and 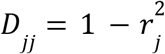 with 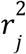 being the imputation quality of variant *j*. A probabilistic graphical illustration of the model is in Figure S9.

Further, current fine-mapping models rely on the relationship *E*[*y*|*X*] = *X*β assuming*X* is measured without error (i.e. not imputed). However, as we observe 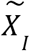 instead of *X*_*I*_ we have 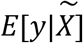 (see Methods):

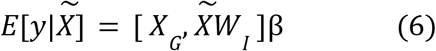

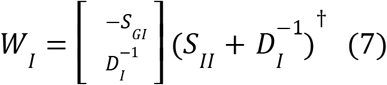

where *S* is the precision matrix of σ, i.e. *S*= σ†. In other words, for an imputed variant, the evidence of being causal depends on both the imputed dosages of the variant and all other variants within the region with their contribution being weighted by *W*_*i*_. As this approach reweights the per-variant evidence based on LD and imputation quality, we refer to this approach as the “reweighting” method. Note that the per-variant likelihood based on the “reweighting” method is similar to a TWAS approach (Gamazon et al. 2015) as both approaches use a weighted sum of variants as the new variable in an association test. By applying the TWAS formula (Gusev et al. 2016; Barbeira et al. 2018), the reweighted statistics can be calculated from GWAS marginal statistics, imputation quality and the LD matrix of a locus from a reference panel without the requirement of individual-level data (Supplementary Notes 2.2).

Applying the “reweighting” scheme followed by fine-mapping, we found that, among experiments with the greatest power (i.e. 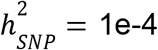 and median p-value is 1.6e-33), the observed coverage of 95% credible sets improved from 89.7% to 91.5% overall and improved from 88.9% to 91.7% if limited to experiments with a common focal variant (MAF > 0.01) (dashed lines in Figure 2 and Figure S5). In cases where the default credible set fails to include the causal variant but captures highly correlated and well imputed neighboring variants, the “reweighting” approach is able to correct such errors (Figure S10). Even though the current error model makes simplifying assumptions such as the independent errors between variants and independent and identically distributed errors among samples, the improved performance over the default approach suggests that our assumptions on the error pattern captures important aspects of the problem and is useful in practice.

### Reweighted credible sets have higher replication rates genome-wide

We next sought to survey the influence of imputation genome-wide using real GWAS that provides a more realistic representation of imputation quality, locus LD, and associations. We leveraged genetic associations based on imputed dosages (Karczewski et al. 2024) (panUKBB) and exome-sequencing (Backman et al. 2021) (ExWAS) from the UK Biobank. As expected, we observed attenuation of marginal statistics in the GWAS relative to those obtained in ExWAS (Figure S11 and Supplementary Notes 2.3).

Next, we evaluated the impact of imputation errors on fine-mapping. We performed fine-mapping using summary statistics-based SuSiE (Zou et al. 2022) on the standing height GWAS in the panUKBB dataset. SuSiE was run with summary statistics as is (“default”) or reweighted summary statistics based on our proposed approach (“reweight”). We considered two credible sets as “shared” if the 95% credible set from one approach captured more than 90% of the cumulative PIP of that from the other approach, otherwise, they were considered “private” to one approach. Among 306 loci with converged SuSiE fits using both approaches, the default approach identified 627 credible sets, and the reweighting approach identified 587. Among these credible sets, 562 credible set pairs were shared (Figure 4A). The default approach generated 65 private signals and the reweighting approach generated 25.

**Figure 4:**
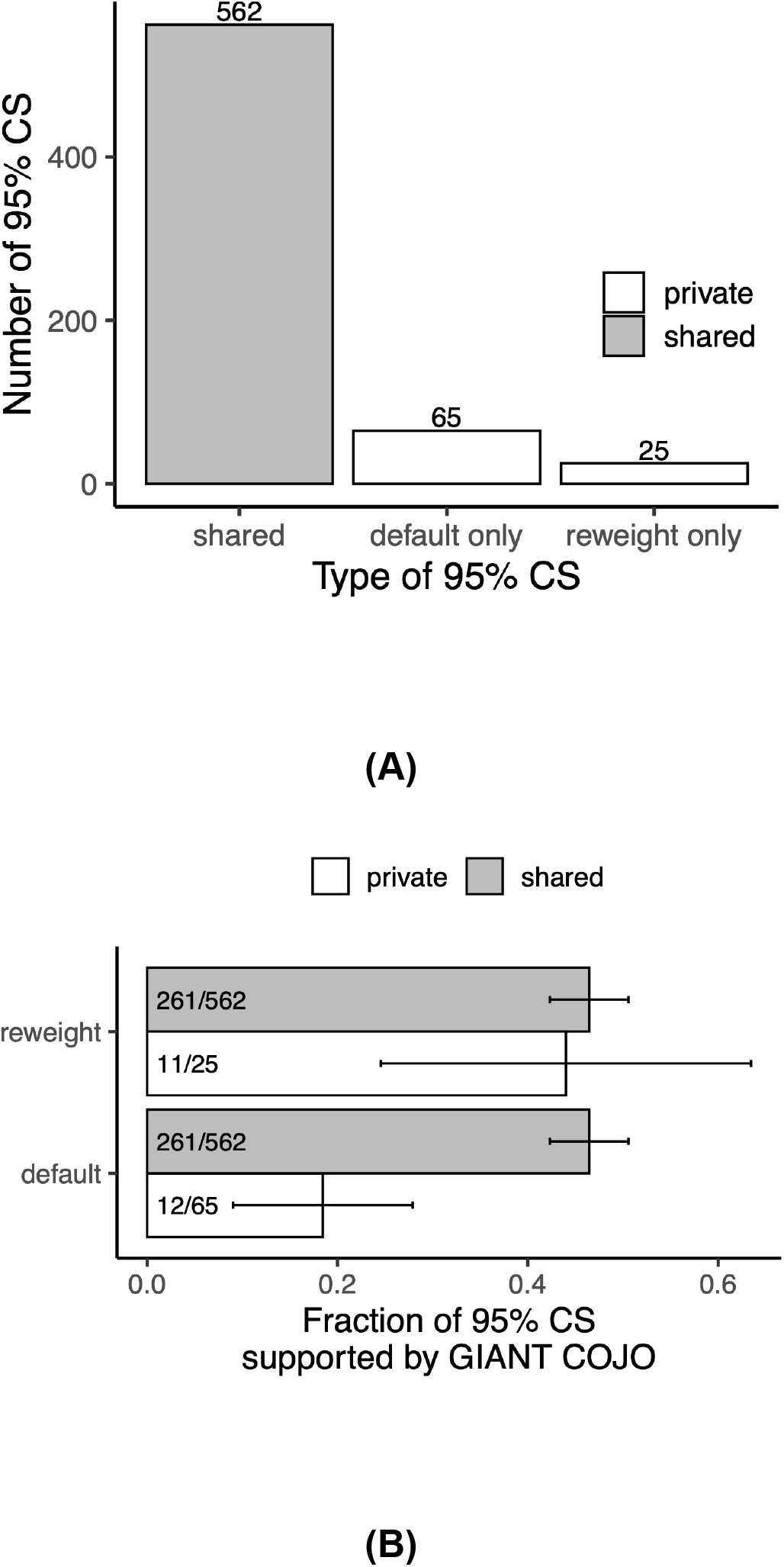
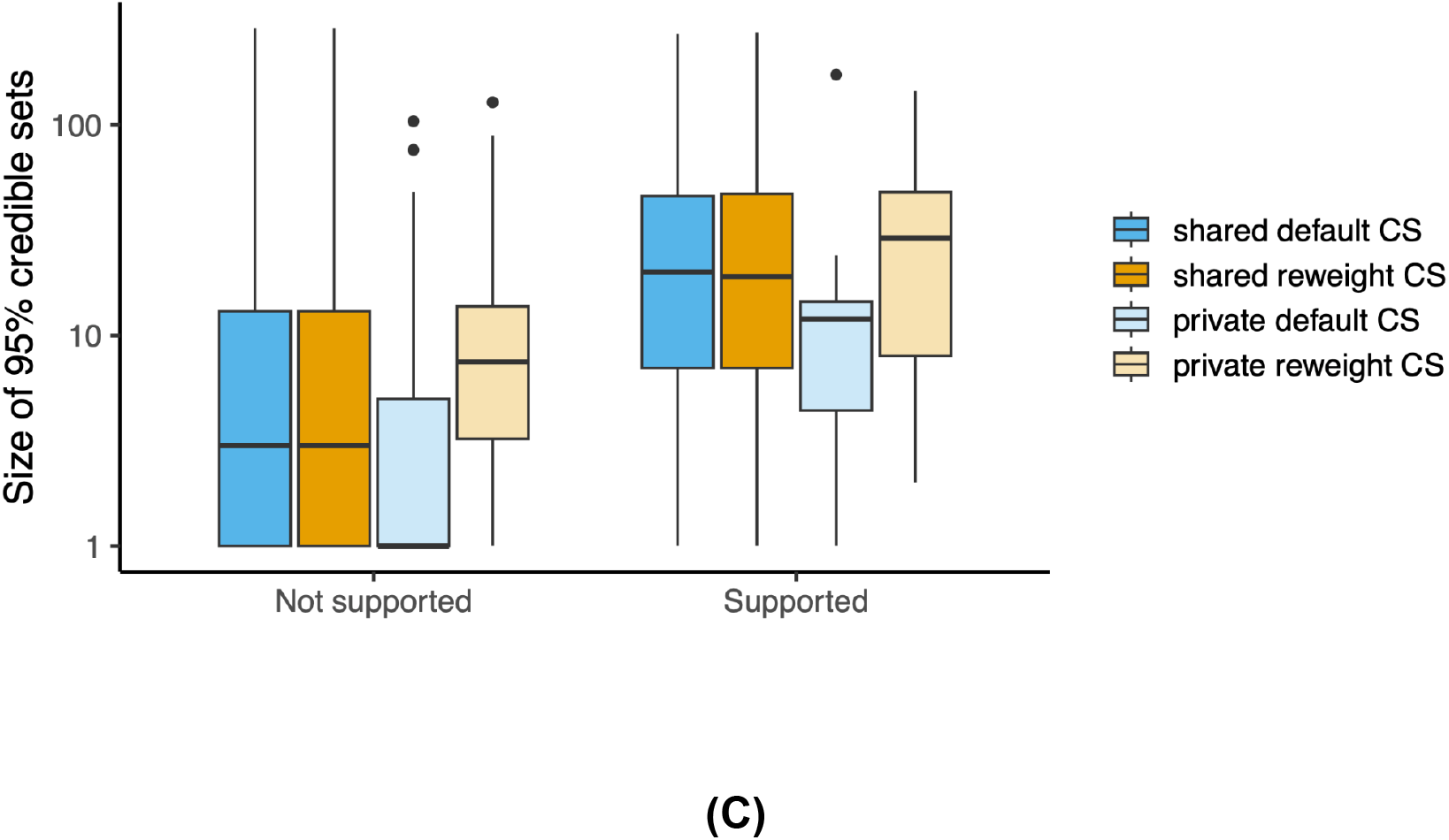
Illustrating the effect of imputation errors in UK Biobank. **(A)** The 95% credible sets by summary statistics-based SuSiE on panUKBB standing height GWAS with default and “reweighting” approaches are classified into three categories: “default only”, “reweight only”, and “shared” based on if the resulting CS is shared between the two approaches. The number of CSs in each category is shown on the y-axis and also as numbers in the plot. The “default only” and “reweight only” CSs are also referred to as “private” CSs which are colored in white. The “shared” CSs are colored in gray. **(B)** For a 95% credible set from either default or “reweighting” approaches, we consider them as being supported if they have maximum *R*^2^ > 0. 90 to any secondary signals in GIANT GWAS within 100,000 bp nearby. The support rate, i.e. the fraction of CSs that are supported, is shown on y-axis for private (in white) and shared (in gray) CSs from default and “reweighting” approaches. The number of supported and all 95% CSs are shown as “#{supported}/#{total}”. The error bar shows the 95% confidence interval. **(C)** The distribution of the size of the 95% credible set is shown on y-axis for credible sets from either default (blue) or “reweighting” (yellow) approaches which are either private (light color) or shared (dark color).

As we had no knowledge of the true causal variants, to quantify the proportion of supported credible sets obtained with and without reweighting, we leveraged signals identified via COJO (Yang et al. 2012) from the largest meta-analysis of height (Yengo et al. 2022) (GIANT GWAS). We used the results from European individuals in the GIANT GWAS, where the UK Biobank contributed 11% of the samples. A 95% credible set was considered to be supported if there existed a signal from the GIANT GWAS *≤* 100,000 bp around the credible set identified via SuSiE and its maximum LD (*R*2) to the variants in the credible set exceeded 0.90. Among the shared 95% credible sets, 46% were supported by the GIANT GWAS (Figure 4B). However, only 18% of the private credible sets from the default approach were supported. In contrast, 44% of the private credible sets obtained from reweighted summary statistics were supported. Similar results were observed when a more stringent criteria was used (R^2^ > 0.95, *≤* 10,000 bp distance) (Figure S12). Among all of the 95% credible sets, the default approach tended to produce smaller 95% credible sets with higher PIPs for top variants compared to the “reweighting” approach regardless of whether the credible set was supported by the GIANT GWAS (Figure 4C and Figure S13). This is consistent with our simulation results, where we observe anti-conservative PIPs when imputation errors exist. The MAFs of the top variant in the credible sets were similar for the default and “reweighting” approaches, which suggested that the differences in the rate of support and size of credible sets were unlikely to be driven by differences in MAF, although the associations found in the GIANT GWAS might be biased towards common variants.

To estimate the fraction of low-quality credible sets due to imputation error, we considered unsupported private credible sets as “false discoveries”. Among all of the fine-mapped credible sets identified by the default approach, 8% are likely false discoveries, whereas this fraction decreases to 2% after “reweighting”. These results suggest the widespread effect of imputation errors on fine-mapping results in real data and show the potential of the “reweighting” approach in reducing false discoveries.

### Reweighted fine-mapping captures known causal variants in a real locus

To illustrate the effect of imputation on fine-mapping through a single locus, we performed fine-mapping on the known F5 gene locus associated with blood clotting disorders using data from 23andMe. The F5 gene encodes coagulation factor V which is one of the proteins that form the coagulation system with several known variants that contribute to clotting disorders. Among the known variants, rs6025 is the well-studied Factor V Leiden variant. In addition, the HR2 haplotype has been reported to contribute to the activated protein C resistance phenotype which results in an increased risk of venous thromboembolism (Bernardi et al. 1997; Faioni 2001). We performed conditional analysis and identified three additional independent associations (conditional step 2 to 4) where the lead variant is within the F5 gene body (p-value < 1e-10) (Supplementary Notes 2.4). To pinpoint the causal variants of the three conditional associations within the F5 gene, we isolated each of the signals and performed the fine-mapping procedure while also applying the “reweighting” approach to model imputation error. In steps 2 and 3, the 95% credible set of both the default and “reweighting” approaches captured the same missense variants (Figure S14). In particular, two HR2 haplotype variants (rs4524 and rs4525) were captured in step 2. The 95% credible set from the “reweighting” approach captured another missense variant in the HR2 haplotype (rs6033) that was missed by the default approach (step 4; Figure 5).

**Figure 5:**
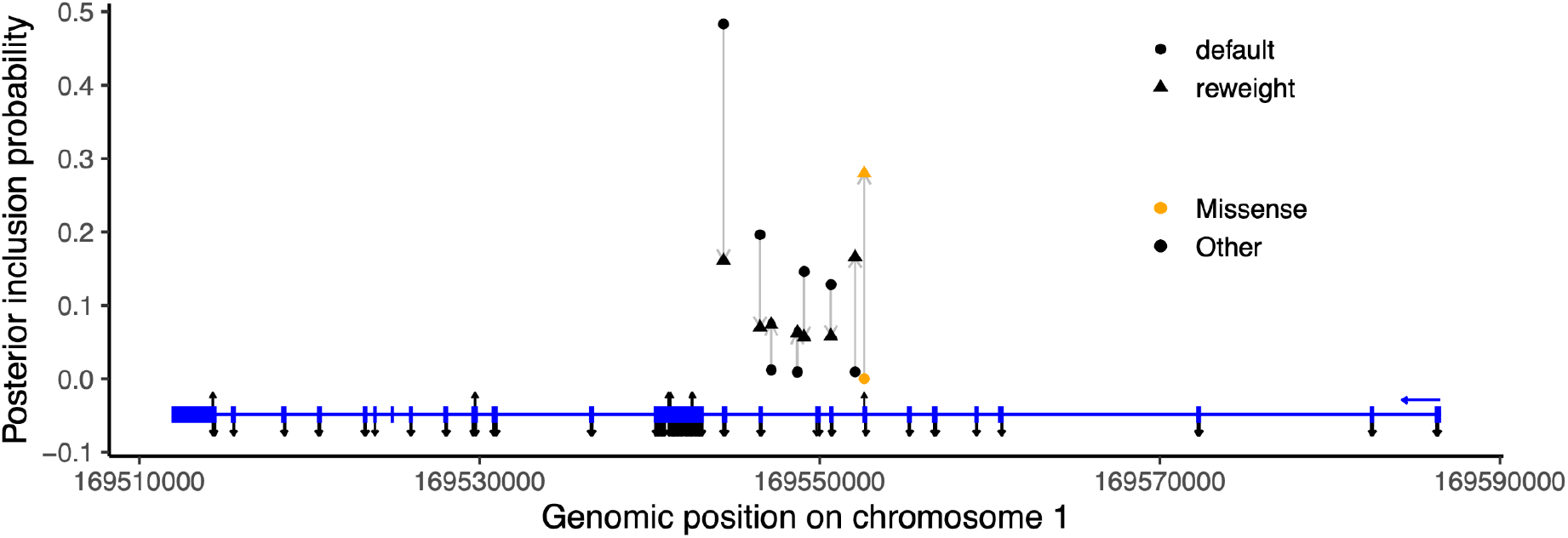
The 95% credible set at the F5 locus. The PIPs of variants in the 95% credible set of the default approach (circles) and the “reweighting” approach (triangles) are shown for the conditional signal step 4 (y-axis). For each variant, the gray arrow points from the PIP of the default approach to the PIP of the “reweighting” approach indicating the direction of change between the two methods. Missense variants are colored in orange and variants without a functional coding consequence are shown in black. The exons of the F5 gene are shown in the bottom in blue with the blue arrow indicating the direction of transcription. HR2 haplotype variants are annotated as upward pointing black arrows among all other missense variants indicated with downward pointing arrows along the F5 exons.

In this locus, the “reweighting” approach had a larger impact on the variants with relatively low imputation quality (Figure S15). However, all candidate fine-mapped variants have *r*^2^ > 0. 94. This further suggests that even considerably high-quality imputation can still affect fine-mapping results. Such influence is highly dependent on the LD pattern of the locus and the power of the GWAS (Supplementary Notes 2.5 and Figure S16).

## 3 Discussion

In this work, we designed a simulation study to look into the effect of genotype imputation on post-GWAS fine-mapping results. First, we quantified the coverage of credible sets and found that the observed coverage was influenced by association strength and the MAF of the true causal variant. Credible sets were miscalibrated when association strength was high and when the true causal variant was in a high LD environment (Figure S5). This was at first a counterintuitive result. However in fine-mapping, imputation accuracy affects statistical power, and in combination with the LD of the locus, both influence the posterior inclusion probabilities from Bayesian fine-mapping (Supplementary Notes 2.5 and Figure S16). For causal variants with many linked non-causal variants, it would require greater statistical power to distinguish the causal variant from those in LD with it. Hence, credible set coverage from fine-mapping analyses should be interpreted with caution especially when the following three conditions are met: i) imputation accuracy is limited; ii) the region has an high LD environment; iii) the association strength is strong.

We also considered the case when the causal variant was absent from the fine-mapping analysis and quantified how well the credible set could tag the missing causal variant. We found that the median LD between the causal variant and the variants in the 95% credible set was much smaller when the causal variant is missing than otherwise regardless of whether the causal variant was genotyped or imputed. Such discrepancy is the largest for rare causal variants. This result highlights the importance of including imputation-based variants in the fine-mapping analysis. This is especially important for a rare causal variant which is usually poorly tagged on a genotyping chip.

On one hand, imputation is detrimental to the ability of a credible set on capturing the causal variant via introducing uncertainties, whereas on the other hand, it improves the ability of a credible set on tagging the causal variant via enlarging the variant set being considered. The common practice of keeping only well imputed variants tries addressing the first part of the problem but misses the opportunity to identify additional signals among less well imputed variants which are majorly rare variants so that their fine-mapping results are less prone to imputation errors. So, to leverage the benefit of imputation while reducing the effect of imputation uncertainty, it is important to develop methods accounting for imputation errors.

To mitigate the effect of genotype uncertainty on fine-mapping outcomes, we proposed a simple reweighting scheme which models imputation quality explicitly. Via simulations, analyzing GWAS from the UK Biobank, and a case study on F5 gene locus for blood clots, we showed improved performance from the “reweighting” method on pinpointing causal variants. This suggested the importance of modeling imputation errors when performing fine-mapping and showed the promise of re-calibrating miscalibrated credible sets.

It should be noted that the conclusions above were based on a very specific study design. In particular, in our simulation study, we considered cases where there was only one causal variant and we only tested one fine-mapping method (Wellcome Trust Case Control Consortium et al. 2012) with a fixed prior across all variants in the locus. We note that the choice of prior does affect the credible set quality when the association strength is low but the impact imputation errors on the credible sets persist regardless of the choice of priors (Figure S17).

Furthermore, unlike previous work which compared results between a locus with fully sequenced variants and that with only imputed variants (van de Bunt et al. 2015), we made the contrast only for the causal variant. In our simulations, we observed 99% coverages for 95% credible sets when genotype values were used and these results cannot be fully explained by a > 0.95 cumulative posterior probability of these credible sets (Figure S18) so the reported credible sets were conservative. However, given we only assessed the effect of the focal variant being genotyped or imputed while keeping all other variants being imputed, this led to conservative PIP estimates for non-focal variants and make the focal variant likely to rank higher than the non-focal ones, which in turn contributed to the overall higher coverage of the credible set. When the focal variant is imputed, the credible set coverage is still conservative for scenarios of small 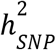 When 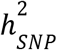 is small, the influence of the competition between imputation quality and the maximum LD nearby on credible set calibration is small (Supplementary Notes 2.5 and Figure S16) and the observed coverage gradually gets lower as the influence of imputation gets amplified by a bigger 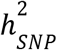. In spite of these limitations, we think the conclusions being drawn are generalizable. Future work is required to examine this topic more comprehensively.

Taken together, we emphasize the importance of having genotype values for causal variants in the GWAS variant set for fine-mapping (via genotyping or imputation), however, we also flag the potential miscalibration of fine-mapping credible sets due to the uncertainty introduced by genotype imputation. To maximize the utility of fine-mapping analyses in the context of a workflow that combines genotyping with imputation, we urge methodological improvements for comprehensively considering imputation error in fine-mapping analyses in the future.

## 4 Methods

### 4.1 Variant selection

We leveraged the 23andMe database for selected participants who were genotyped on a custom SNP array based on the Illumina Global Screening Array, consisting of approximately 654,000 preselected variants and approximately 50,000 custom content variants. Participants provided informed consent and volunteered to participate in the research online, under a protocol approved by the external AAHRPP-accredited IRB, Ethical & Independent (E&I) Review Services. As of 2022, E&I Review Services is part of Salus IRB (https://www.versiticlinicaltrials.org/salusirb).

We selected variants with both genotyped and imputed values. To do so, we started with the set of genotyped variants which were excluded to be part of the imputation scaffold. Among these variants, to keep only high-quality genotyped variants, we further filtered by two genotyping quality criteria: i) genotyping rate > 0.95, and ii) the genotyping quality (measured by the squared correlation between the genotyped values and additional WGS based values as ground truth) > 0.9 among at least 1000 observations.

In total, 9,509 variants passed the filters above. We downsampled these variants by MAF and imputation *r*2(the average value across batches) bins. Specifically, MAF bins are (1e-6, 1e-4], (1e-4, 1e-3], (1e-3, 1e-2], (1e-2, 5e-2], (5e-2, 0.1], (0.1, 0.2], (0.2, 0.3], (0.3, 0.4], and (0.4, 0.5]. The imputation *r*2 bins are (0.3, 0.4], (0.4, 0.5], (0.5, 0.6], (0.6, 0.7], (0.7, 0.8], (0.8, 0.9], and (0.9, 1]. For each combination of MAF and imputation *r*2 bins, we randomly selected 10 variants if there were more than 10 and selected all variants otherwise. 402 variants were kept after this downsampling approach.

### 4.2 Simulation procedure

The simulation was performed at locus level where each of the selected variants was treated as the single causal variant of the locus which was called the focal variant. Our simulation included 2 million samples. We simulated phenotypes as *Y* = *X* + ϵ, ϵ *∼ N*(0, σ^2^) where *X* is the genotyped values of the focal variant. And thecausal effect size is always 1 in our simulation. σ^2^ was set based on the desired locus-level heritability (i.e. per-SNP heritability,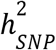). Roughly, the expected *χ*^2^ statistics of the association test is approximately 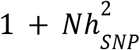 with *N* being the sample size. As we used 2 million samples for the simulation study, to cover a wide range of statistical power, we set 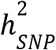 to be 5e-6, 1e-5, 1.5e-5, 2.5e-5, 5e-5, and 1e-4. These correspond to having the association p-value centering around 1.6e-3, 7.7e-6, 4.3e-8, 1.5e-12, 1.5e-23, and 2.1e-45.

### 4.3 Fine-mapping procedure

For each of focal variants, all genotyped and imputed variants within 100kb up/downstream were considered and among these we further removed variants with imputation *r*^2^< 0.3 or differential values between batches or genotyping rate < 0.9.

For each locus, we first ran the association test for all variants being included to obtain the marginal test statistics. Then, we performed fine-mapping analysis as proposed by (Wellcome Trust Case Control Consortium et al. 2012) using these summary statistics. The prior parameter *W*, the variance of the effect size, was set to 0.1. The fine-mapping was ran with three settings, in which the summary statistics of the non-focal variants were always included but, for the focal variant, we included: i) summary statistics from the genotyped values; ii) summary statistics from the imputed values; iii) no summary statistics of the focal variant. These three settings correspond to the cases that: i) the true genotypes of the causal variant are observed (ideal case); ii) the causal variant is imputed with error; iii) the causal variant is missing.

### 4.4 Deriving “reweighting” formula

Recall that to consider imputation error explicitly, we propose the following model (Equations 3, 4, 5 in main text)

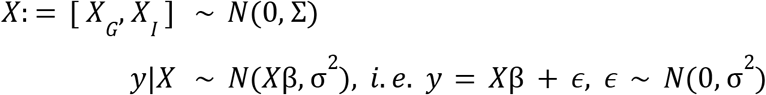

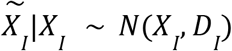

where *y* is the phenotype, *X* is the actual genotype, and 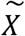 is the imputed dosage with *D*_*I*_ representing the imputation error. The subscripts *G* and *I* represent the blocks of genotyped and imputed variants separately.

Fine-mapping methods rely on *E*(*y*|*X*) = *X*β. However, as we don’t observe 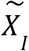, we want to derive 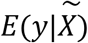 instead.

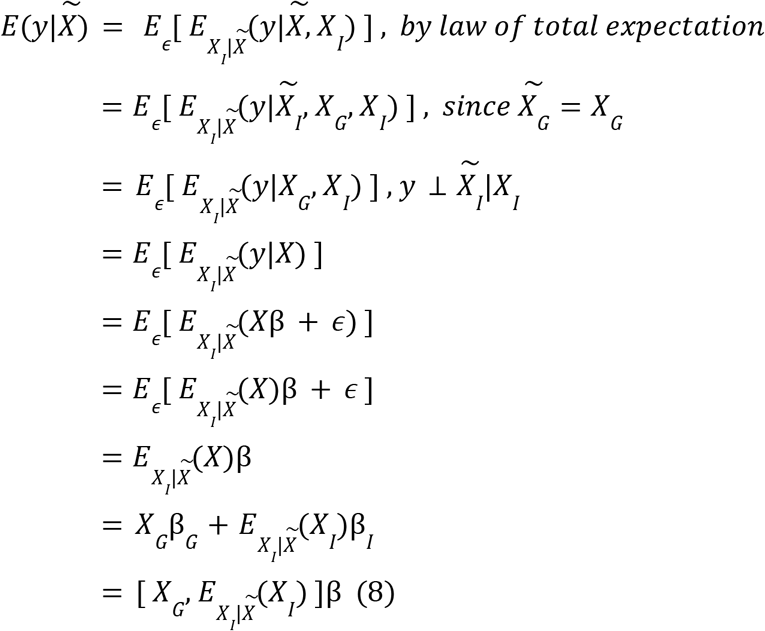

To derive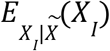, consider 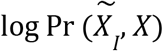.

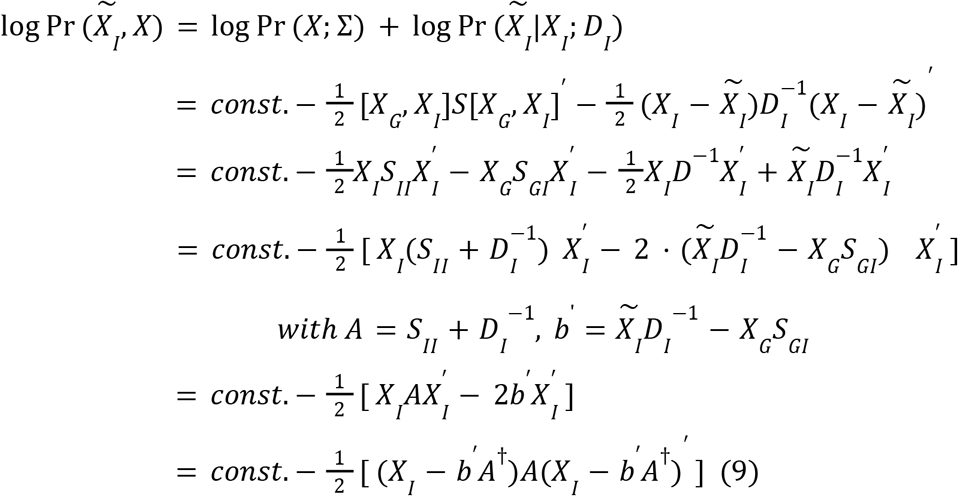

Where *S* = σ^†^. Here *const*.includes all terms that are not related to *X*_*I*_. Based on Equation 9, we have

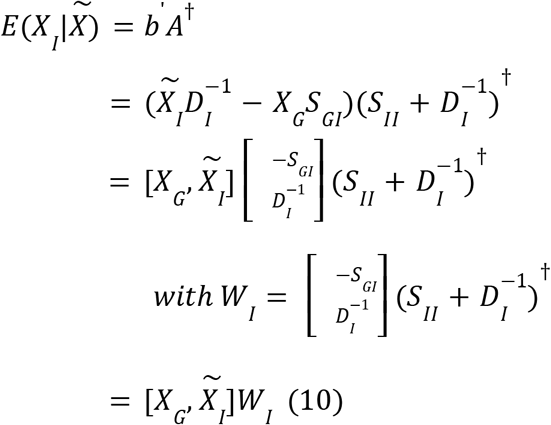

Taking together Equations 8 and 10

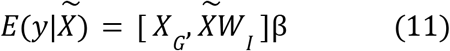

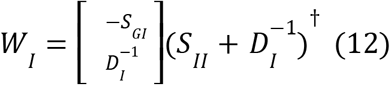

## Acknowledgements

We thank Dr. Jared O’Connell for help and advice on the variant selection for the simulation study. We thank Dr. David A. Hinds for critical review. We would like to thank the research participants and employees of 23andMe for making this work possible.

The following members of the 23andMe Research Team contributed to this study: Stella Aslibekyan, Adam Auton, Elizabeth Babalola, Robert K. Bell, Jessica Bielenberg, Ninad S. Chaudhary, Zayn Cochinwala, Sayantan Das, Emily DelloRusso, Payam Dibaeinia, Sarah L. Elson, Nicholas Eriksson, Chris Eijsbouts, Teresa Filshtein, Pierre Fontanillas, Davide Foletti, Will Freyman, Zach Fuller, Julie M. Granka, Chris German, Éadaoin Harney, Alejandro Hernandez, Barry Hicks, David A. Hinds, Michael V. Holmes, M. Reza Jabalameli, Ethan M. Jewett, Yunxuan Jiang, Sotiris Karagounis, Lucy Kaufmann, Matt Kmiecik, Katelyn Kukar, Alan Kwong, Keng-Han Lin, Yanyu Liang, Bianca A. Llamas, Aly Khan, Steven J. Micheletti, Matthew H. McIntyre, Meghan E. Moreno, Priyanka Nandakumar, Dominique T. Nguyen, Jared O’Connell, Steve Pitts, G. David Poznik, Alexandra Reynoso, Shubham Saini, Morgan Schumacher, Leah Selcer, Anjali J. Shastri, Jingchunzi Shi, Suyash Shringarpure, Keaton Stagaman, Teague Sterling, Qiaojuan Jane Su, Joyce Y. Tung, Susana A. Tat, Vinh Tran, Xin Wang, Wei Wang, Catherine H. Weldon, Amy L. Williams, Peter Wilton.

