## Supplementary Materials for "Credible set is sensitive to imputation quality and missing variants"

### 1 Supplementary Figures

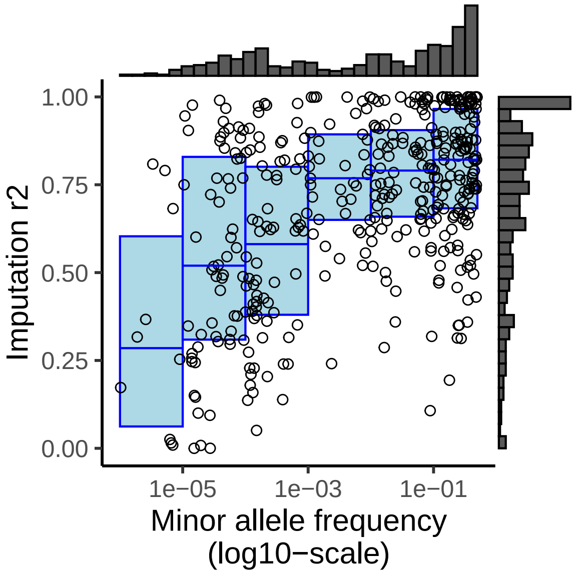

Figure S1: **Imputation quality and minor allele frequency of focal variants.** For all 402 focal variants being included in the simulation study, a scatter plot of imputation quality (y-axis) versus minor allele frequency (x-axis) is shown. The imputation quality is defined as the squared correlation between genotyped and imputed values. The histograms of the imputation quality and the minor allele frequency are shown on the right and top side of the scatter plot. The blue rectangles represent the 25% and 75% quartiles of the imputation quality for a given MAF bin with a horizontal line in the middle representing the median of the imputation quanlity. Six rectangles correspond to MAF bins: (1e-06,1e-05], (1e-05,0.0001], (0.0001, 0.001], (0.001, 0.01], (0.01,0.1], (0.1,0.5].

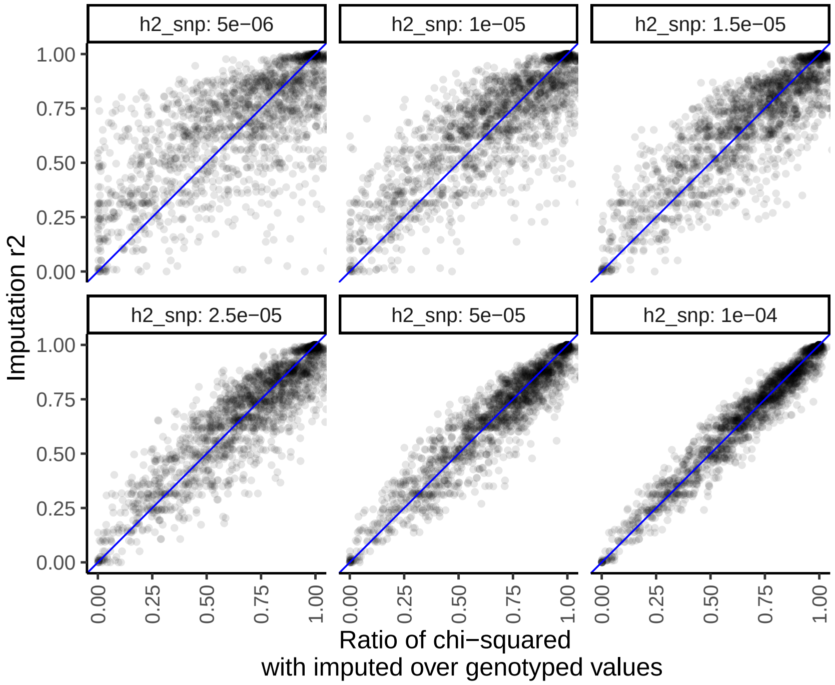

Figure S2: **Ratio of GWAS $\chi^2$ for results from imputed over genotyped values.** For each of the 402 focal variants, the ratio of the marginal $\chi^2$, $\chi^2_{\text{imputed}}/\chi^2_{\text{genotyped}}$, is shown on x-axis and the corresponding imputation quality of the imputed values is shown on y-axis. Each panel corresponds to a $h^2_{\text{SNP}}$ value in the simulation. The blue line is the $y = x$ line.

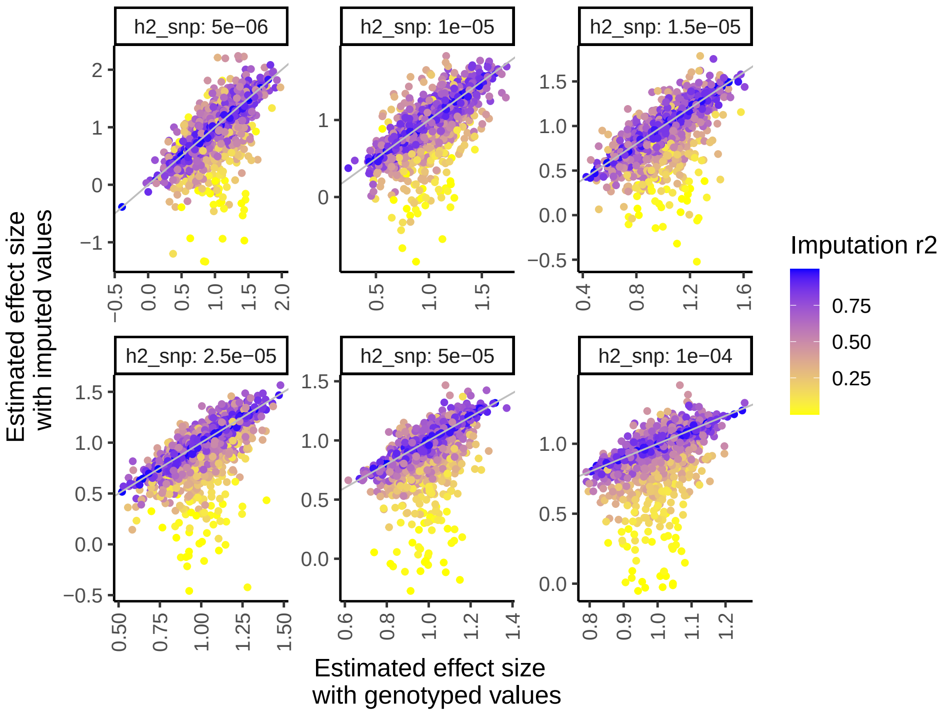

Figure S3: **Comparing marginal estimated effect sizes between genotyped and imputed values.** For each of the 402 focal variants, the marginal estimated effect sizes obtained by using genotyped (x-axis) and imputed values (y-axis) are shown as scatter plots. Each panel corresponds to a $h^2_{\text{SNP}}$ value in the simulation. The color indicate the imputation quality of the imputed values. The gray line is the $y = x$ line.

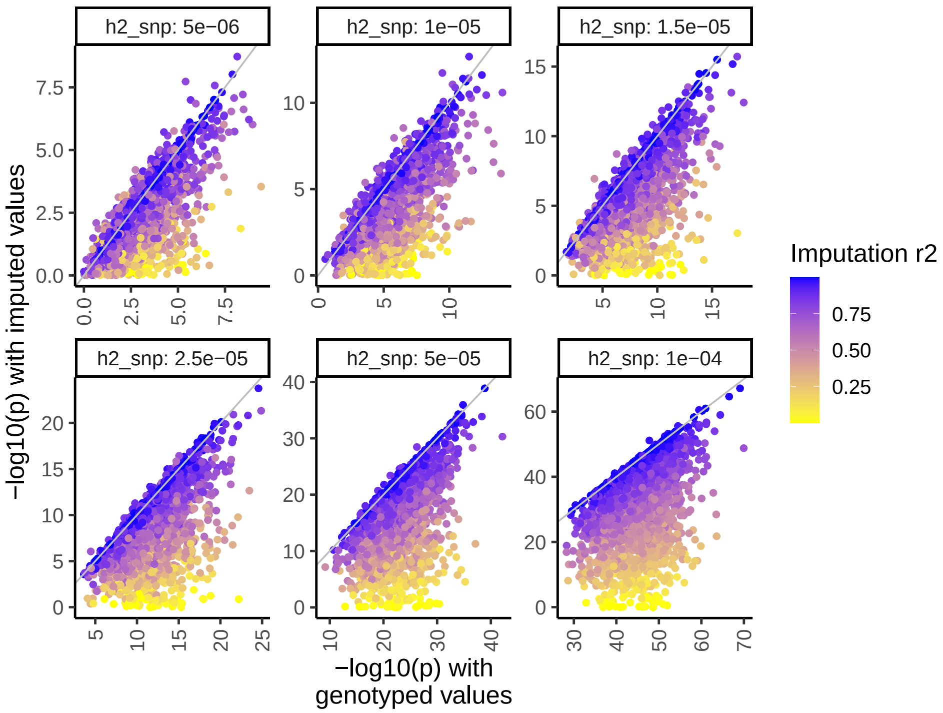

Figure S4: **Comparing marginal $-\log_{10}(p)$ between genotyped and imputed values.** For each of the 402 focal variants, the marginal $-\log_{10}(p)$ obtained by using genotyped (x-axis) and imputed values (y-axis) are shown as scatter plots. Each panel corresponds to a $h^2_{\text{SNP}}$ value in the simulation. The color indicate the imputation quality of the imputed values. The gray line is the $y = x$ line.

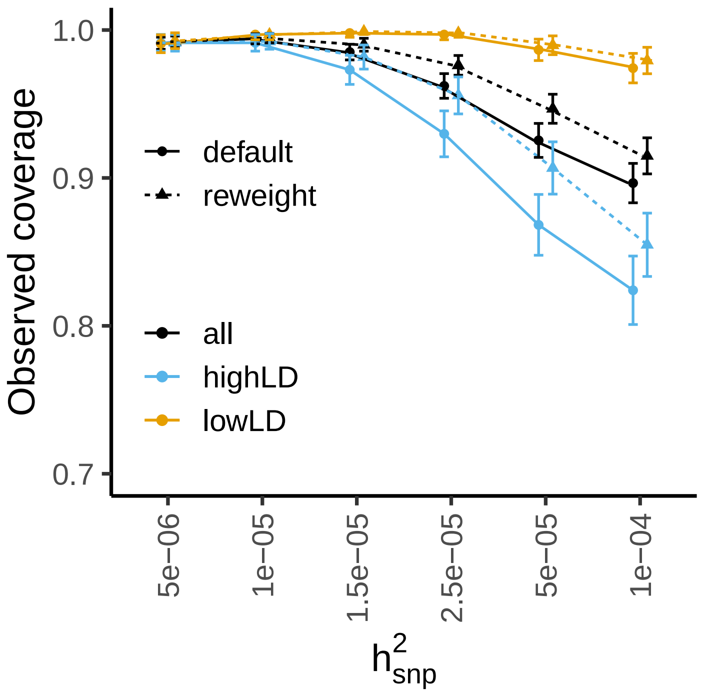

Figure S5: **The observed coverage of 95% credible sets with imputed genotypes stratified by LD environment.** At each $h^2_{\text{SNP}}$ (x-axis), the observed coverage is shown (y-axis) stratified by LD environment of the focal variant which is defined as the maximum LD $R^2$ between the focal variant and any other variants within the locus (±100kb) (highLD: max $R^2 > 0.5$; lowLD: max $R^2 < 0.5$). The results based on imputed genotypes are shown. Solid lines show the results based on the association result as-is. Dashed lines show the results based on the proposed "reweighting" approach.

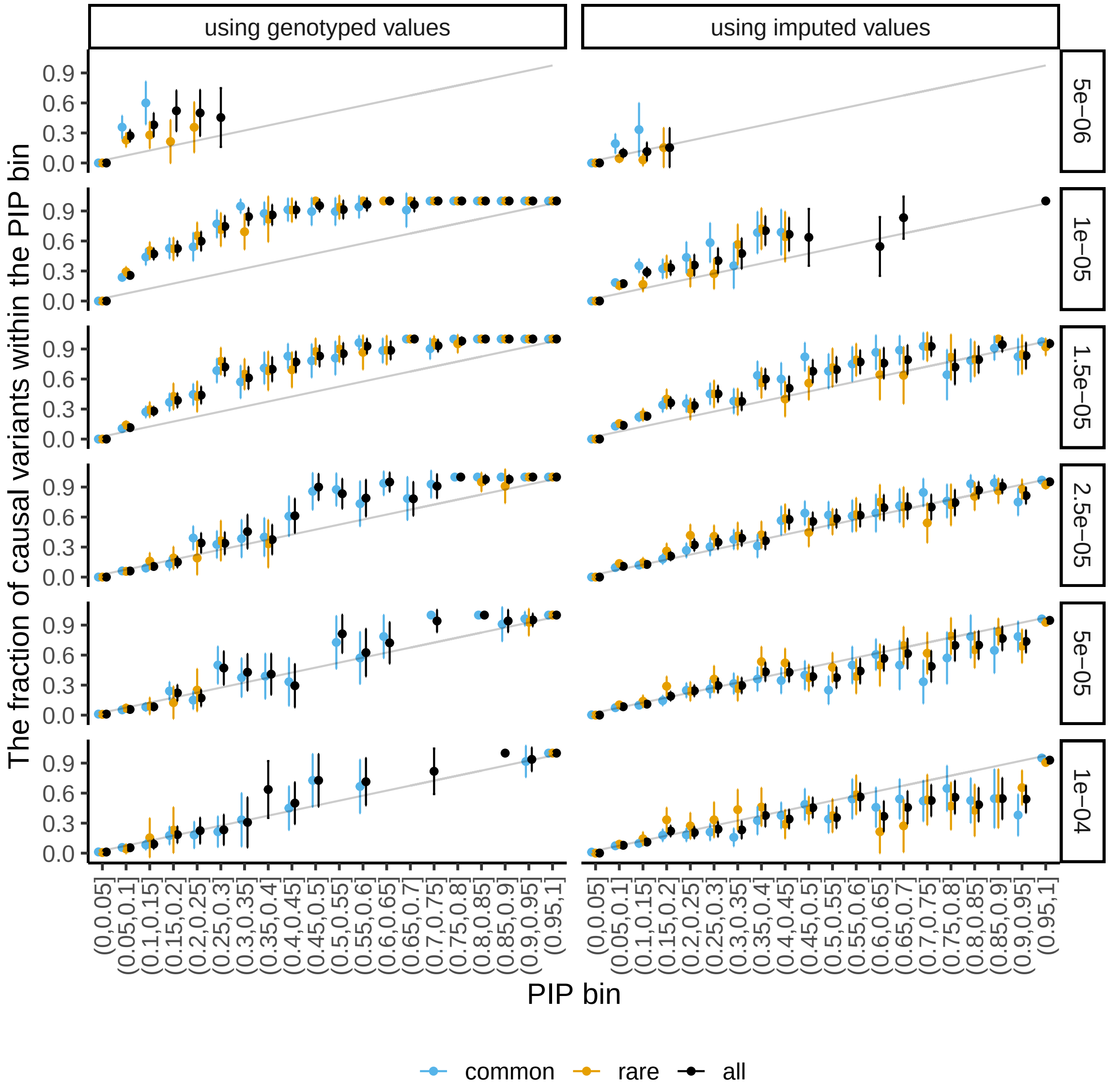

Figure S6: **Examining the calibration of the posterior inclusion probability stratified by MAF.** We examined the calibration of the posterior inclusion probability for each experimental setup, i.e. for a given $h^2_{\text{SNP}}$ value (panels organized in rows) with focal variants being either genotyped or imputed (panels organized in columns). Across all experiments for an experimental setup, all variants are stratified by their posterior inclusion probabilities in 20 equally-spaced bins between 0 and 1 which we called PIP bins. Within each PIP bin, the fraction of causal variants is shown on y-axis and the error bar indicates the standard error of the estimated fraction. The bin with fewer than 10 variants are not shown. The gray line corresponds to the expected fraction of causal variants when posterior inclusion probablity is perfectly calibrated (PIP is uniformly distributed within the bin), i.e. the mean of the low and high boundary of the PIP bin. To examine the effect of minor allele frequency of the causal variant, the analysis is further stratified by the MAF of focal variants with MAF $< 0.01$ shown in yellow and MAF $> 0.01$ shown in blue. The results without MAF stratification are shown in black.

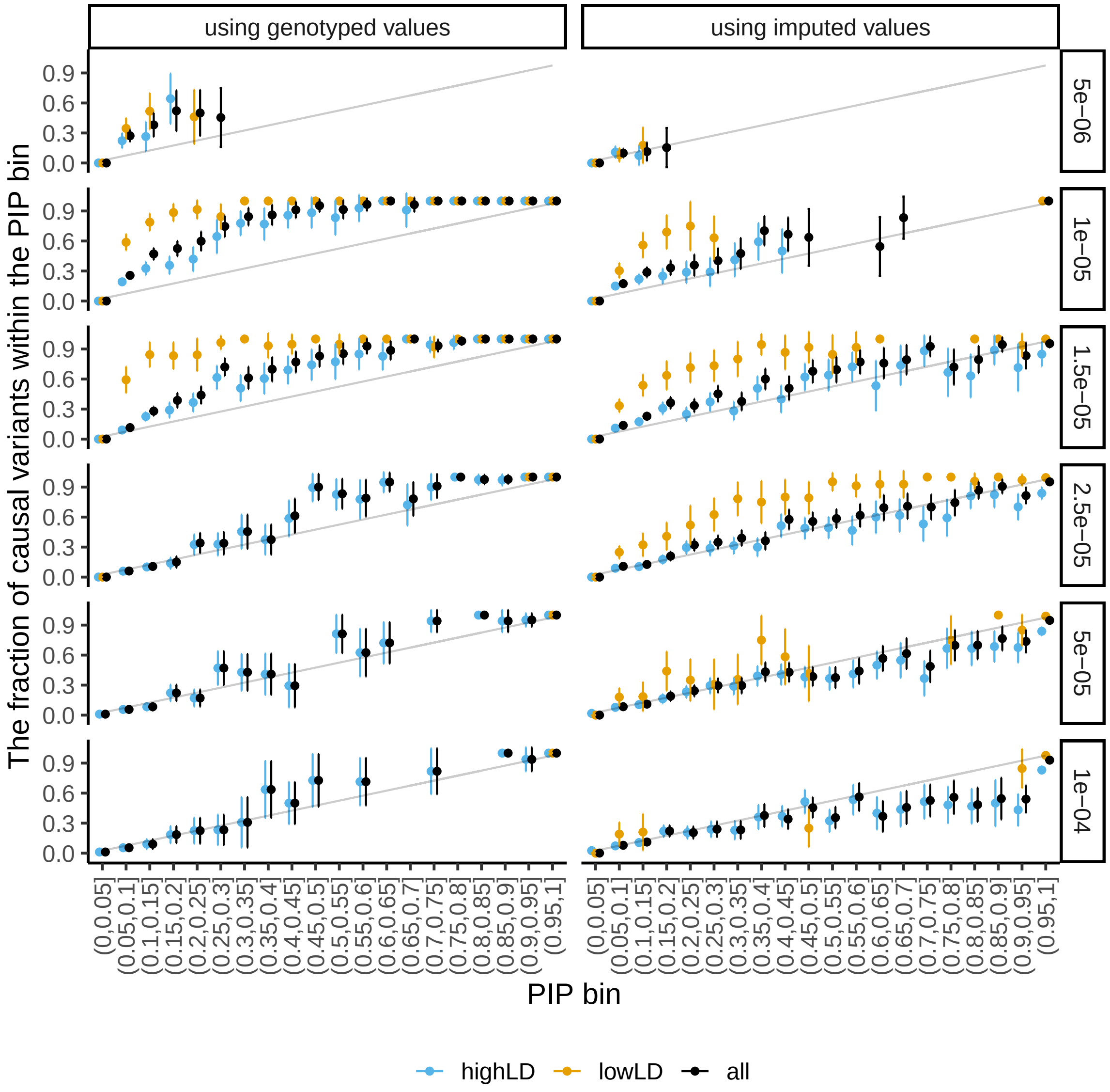

Figure S7: **Examining the calibration of the posterior inclusion probability stratified by LD environment.** We examined the calibration of the posterior inclusion probability for each experimental setup, i.e. for a given $h^2_{\text{SNP}}$ value (panels organized in rows) with focal variants being either genotyped or imputed (panels organized in columns). Across all experiments for an experimental setup, all variants are stratified by their posterior inclusion probabilities in 20 equally-spaced bins between 0 and 1 which we called PIP bins. Within each PIP bin, the fraction of causal variants is shown on y-axis and the error bar indicates the standard error of the estimated fraction. The bin with fewer than 10 variants are not shown. The gray line corresponds to the expected fraction of causal variants when posterior inclusion probablity is perfectly calibrated (PIP is uniformly distributed within the bin), i.e. the mean of the low and high boundary of the PIP bin. To examine the effect of the LD environment of the causal variant, the analysis is further stratified by the maximum LD between the focal variant and any other variants within the locus (±100kb) with highLD - max $R^2 > 0.5$ - shown in blue and lowLD - max $R^2 < 0.5$) shown in yellow. The results without the LD environment stratification are shown in black.

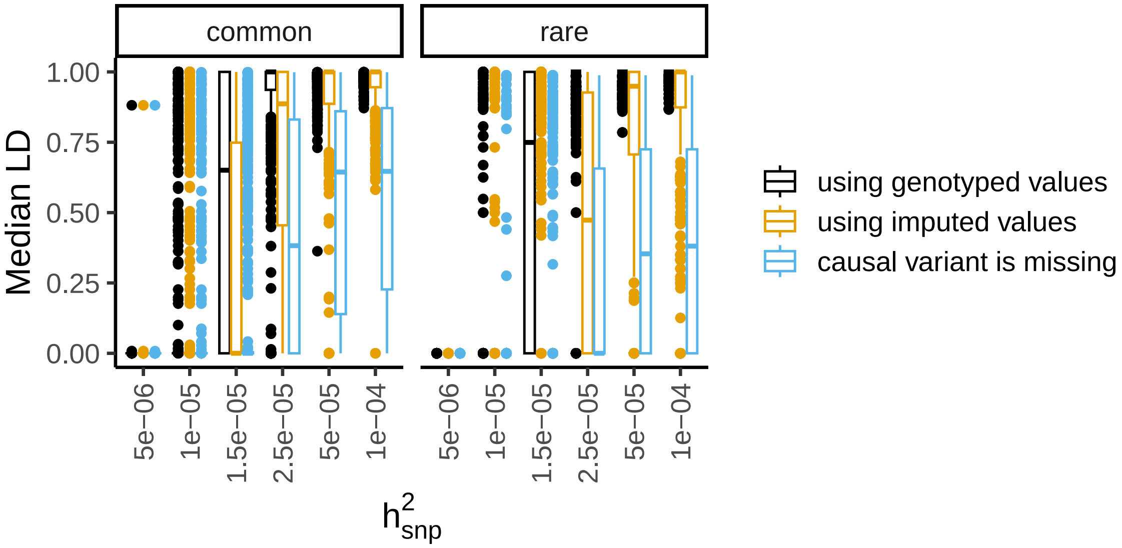

Figure S8: **Median LD $R^2$ between the 95% credible set and the causal variant.** For each experiment, fine-mapping analysis is ran within the locus with: i) genotyped values at the focal variant (black); ii) imputed values at the focal variant (yellow); iii) the focal variant being missing (blue). To measure the quality of the 95% credible set, the median of LD (in terms of the squared correlation) between all the variants among the 95% credible set and the focal variant is calculated. The distribution of the median LD (on y-axis) is shown for each $h^2_{\mathrm{SNP}}$ of the corresponding experiments (on x-axis) and the results for common focal variants (MAF $> 0.01$) and rare focal variants (MAF $< 0.01$) are shown separately.

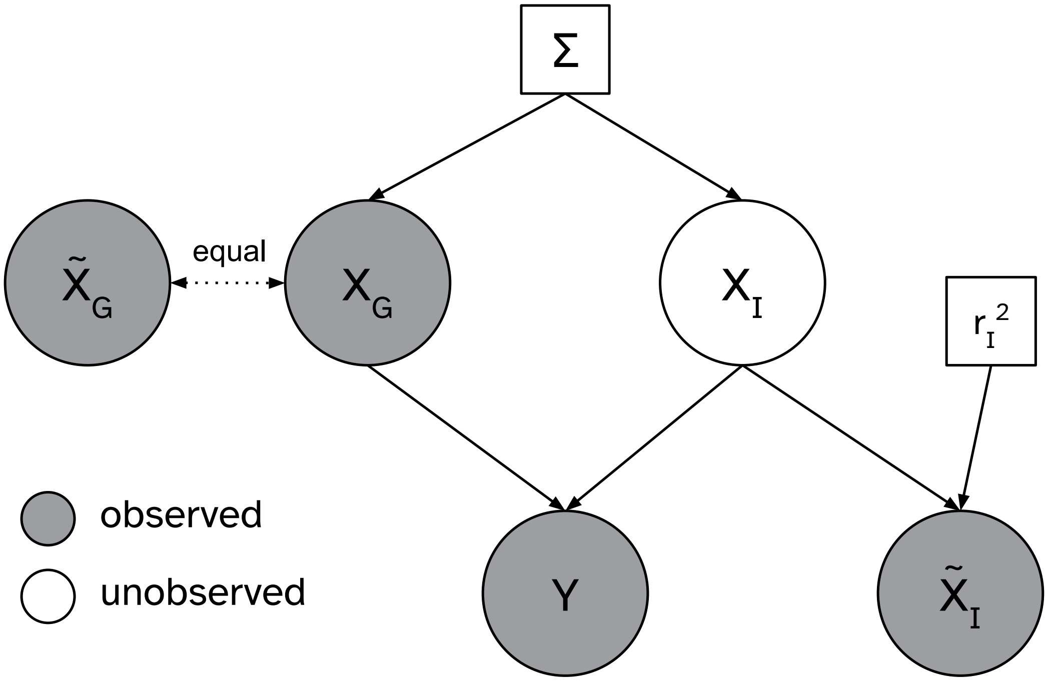

Figure S9: **A probabilistic graphical model representation of the proposed model.** The observed variables are gray nodes and the unobserved variables are white nodes. The parameters of the model are represented by squares and the random variables are represented by circles. $Y$ represents the phenotype and $X$ represents the genotypes with the subscripts $G$ and $I$ indexing the set of variants being genotyped and imputed. $\tilde{X}$ represents the imputed dosages so $X_G$ and $\tilde{X}_G$ are equal which is indicated by dotted line. $\Sigma$ is the LD of the locus and $r_I^2$ is the imputation quality of the imputed dosages. $\beta$ is the causal effect size.

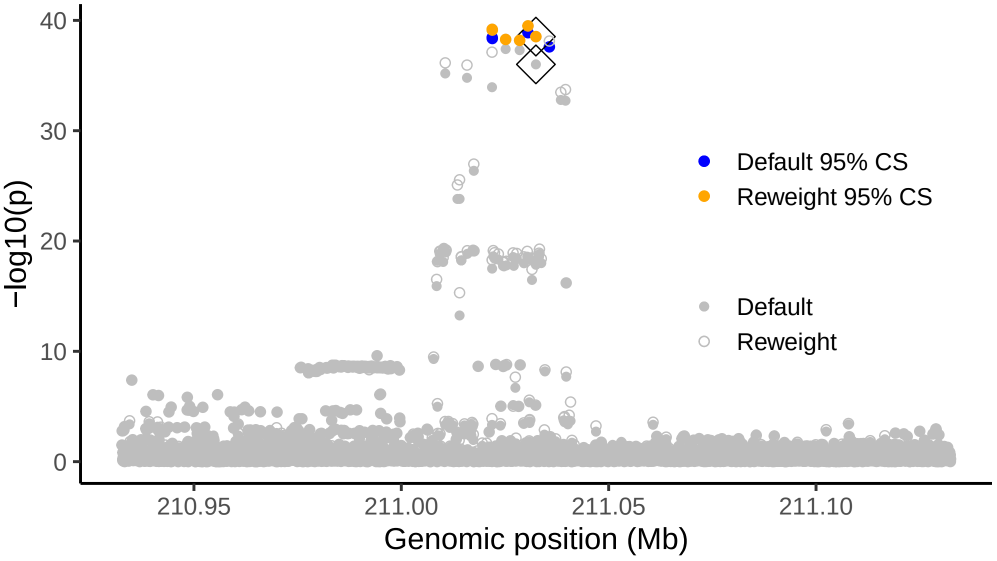

**(A)**

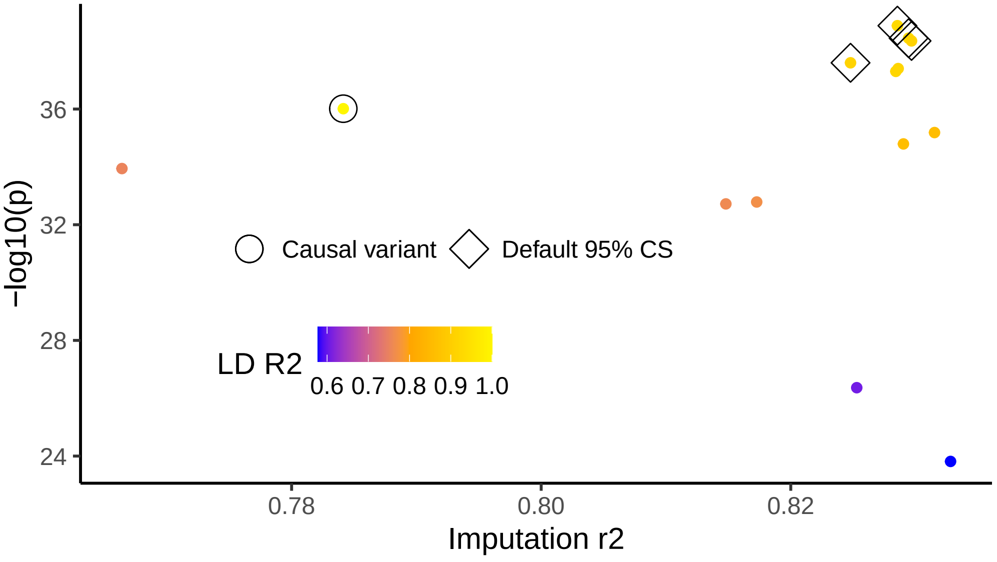

**(B)**

Figure S10: **Illustrating default and the "reweighting" approach in an example locus.**. The simulation results of an example locus on chromosome 2 are shown. **(A)** -log10 p-value of the marginal test and the one after reweighting are shown in gray. The 95% CSs are highlighted in blue (default method) and orange ("reweighting" method). The results of the true causal variant are further labelled by $\Diamond$. **(B)** Imputation r2 (x-axis) vs -log10 p-value (y-axis) for all variants with LD $R^2 > 0.5$ with the causal variant are shown. Color indicates the LD R2. The true causal variant and the 95% CS of the default method are further labelled by $\Diamond$ and $\circ$.

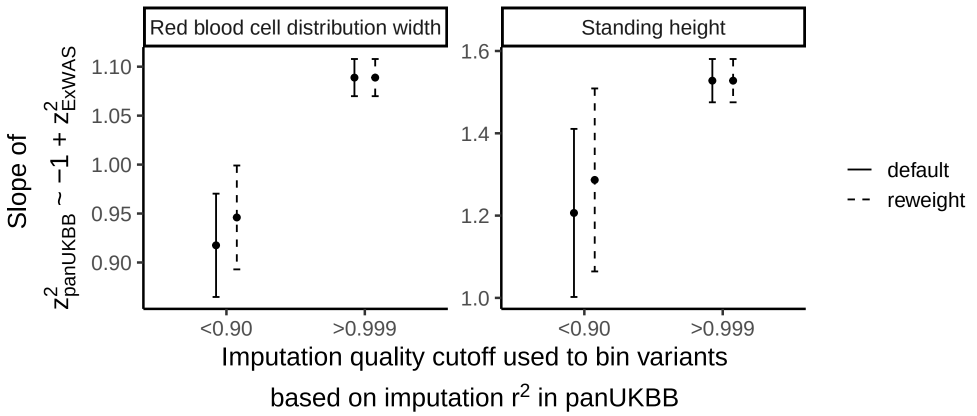

Figure S11: **The effect of imputation errors on GWAS $\chi^2$ in UK Biobank.** Among variants with $p_{\text{ExWAS}} < 0.05$, the slopes of $\chi^2_{\text{panUKBB}} \sim \chi^2_{\text{ExWAS}}$ (via robust regression) are shown on y-axis for two variant sets based on the imputation quality in panUKBB GWAS on x-axis. The intervals present the 95% confidence interval of the estimated slope.

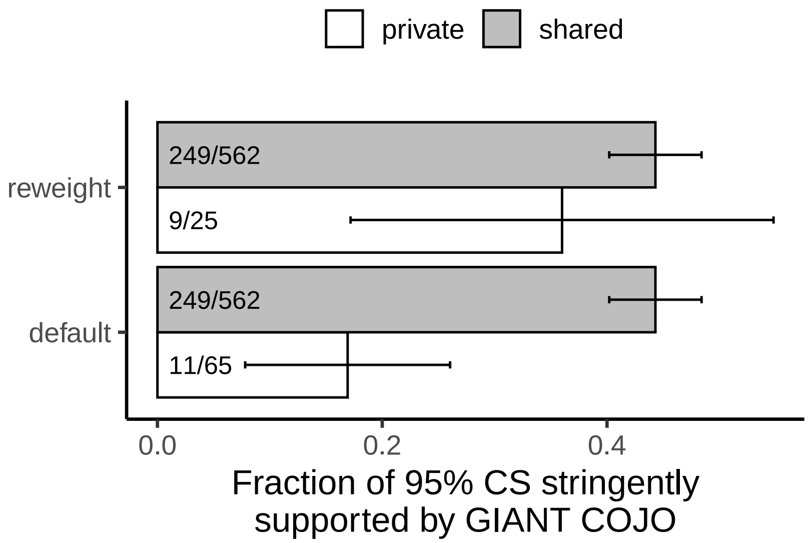

Figure S12: **The fraction of panUKBB 95% credible sets that are stringently supported by GIANT GWAS.** For a 95% credible set from either default or "reweighting" approaches, we consider them as being stringently supported if they have maximum $R^2 > 0.95$ any secondary signals in GIANT GWAS within 10,000 bp nearby. The support rate, i.e. the fraction of CSs that are stringently supported, is shown on y-axis for private (in white) and shared (in gray) CSs from default and "reweighting" approaches. The error bar shows the 95% confidence interval.

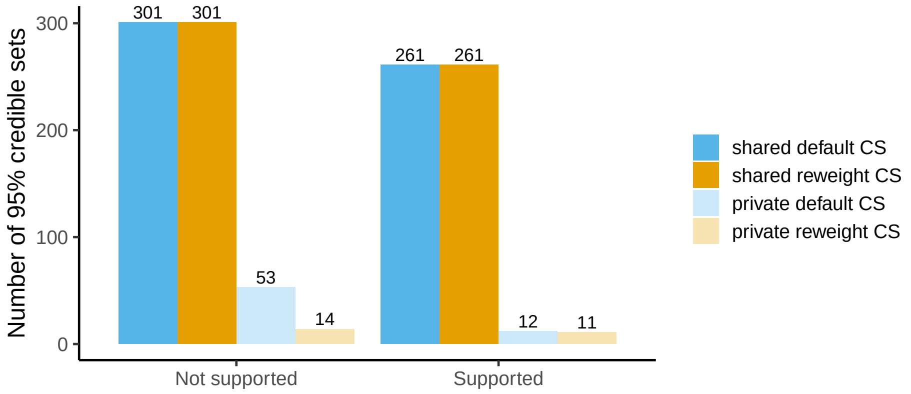

**(A)**

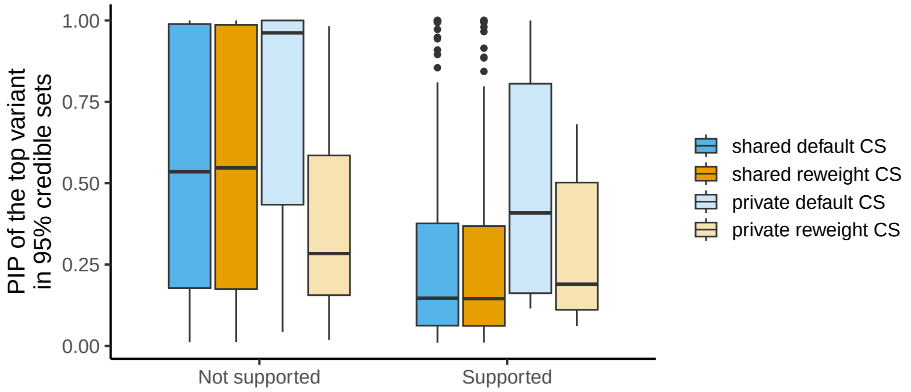

**(B)**

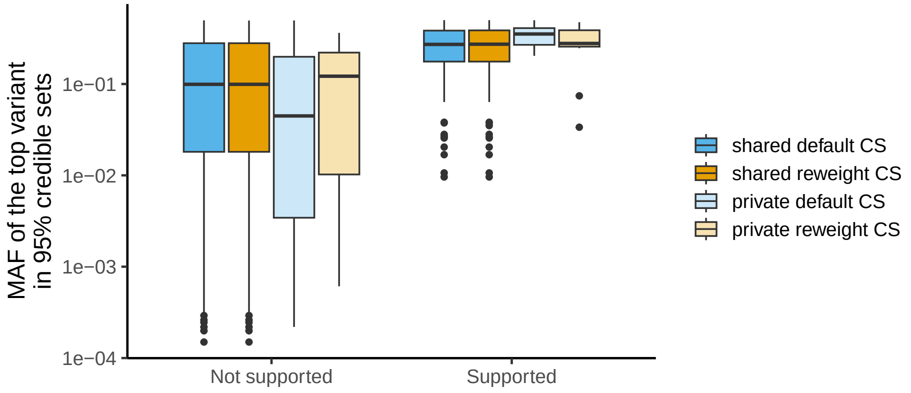

**(C)**

Figure S13: **Characteristics of the 95% credible set identified in panUKBB height GWAS.** Some characteristics of the 95% credible set are shown in which the credible sets are stratified by whether it is classified by a private (light color) or a matched (dark color) credible set and whether the credible set is from default (blue) or "reweighting" (yellow) approaches. **(A)** The number of 95% credible sets falling into each stratum is shown on y-axis. **(B)** The distribution of the PIP of the top variants of the 95% credible set is shown as boxplots on y-axis. **(C)** The distribution of the MAF of the top variants of the 95% credible set is shown as boxplots on y-axis.

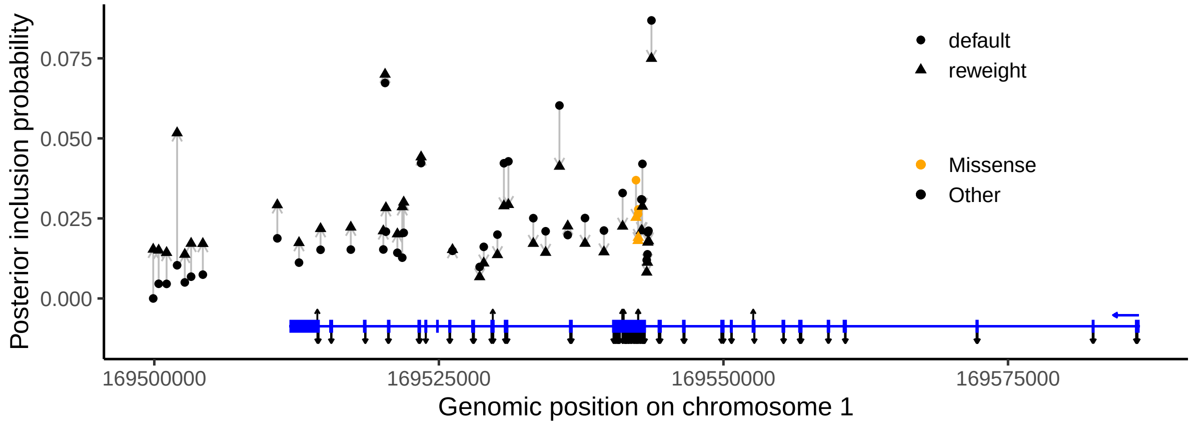

**(A)**

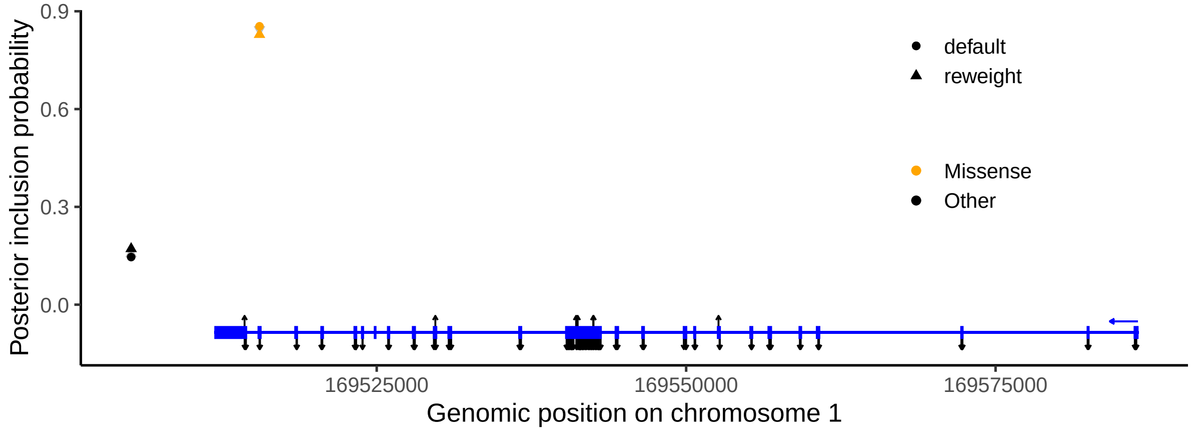

**(B)**

Figure S14: **The 95% credible set of additional two independent signals at the F5 locus.** The PIPs of variants in the 95% credible set of the default approach (circles) and the "reweighting" approach (triangles) are shown for the conditional signal step 2 (panel **(A)**) and 3 (panel **(B)**) on y-axis. For each variant, the gray arrow goes from the PIP from the default approach to the one from the "reweighting" approach indicating the direction of the change between the two methods. The missense variants are colored in orange and variants without a functional coding consequence are shown in black. The exons of the F5 gene are shown in the bottom in blue and the blue arrow on the right indicate the direction of the transcription. The black arrows along the F5 gene exons are missense variants. HP variants point upward and all other missense variants point downward.

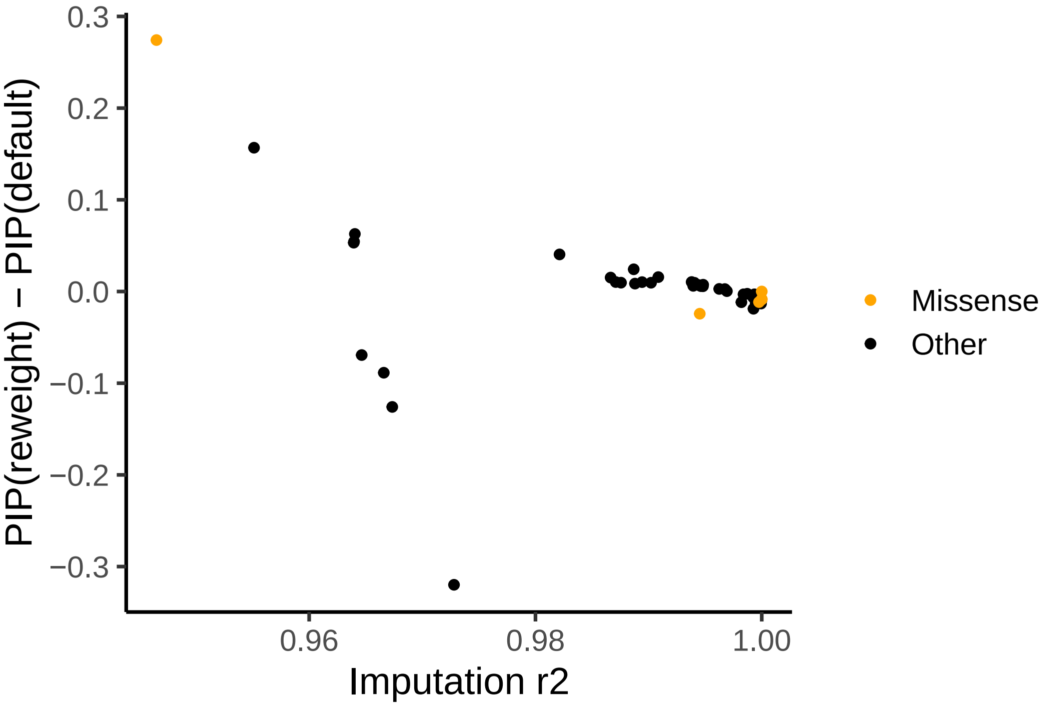

Figure S15: **The difference of PIP between default and the "reweighting" approach versus the imputation quality for the F5 locus of blood clots GWAS.** For all variants within 95% CS, the difference of PIP between default and the "reweighting" approach is shown on x-axis and the imputation $r^2$ is shown on the y-axis. The missense variants are colorred in orange and variants without a functional coding consequence are shown in black.

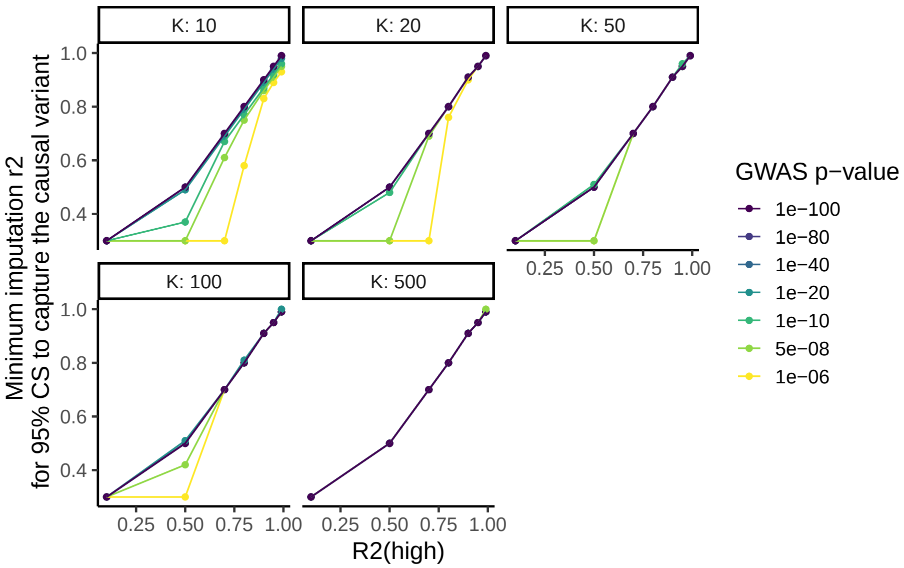

Figure S16: **Dissecting the additional factors determining the effect of imputation quality on fine-mapping.** We constructed an example locus with 2000 variants with various haplotype lengths ($K$, the number of variants within the haplotype), LD environment strength ($R^2$), and GWAS powers (p-values vary from $10^{-6}$ to $10^{-100}$). See details in Supplementary Notes 2.5. We calculated the minimum imputation $r^2$ required in order for the fine-mapping 95% credible set to capture the causal variant at each $K$, $R^2$, and GWAS p-value. The minimum imputation $r^2$ is shown on y-axis as a function of $R^2$ with the colors indicating the different GWAS p-values at the causal variant. Each panel shows the results for a given $K$.

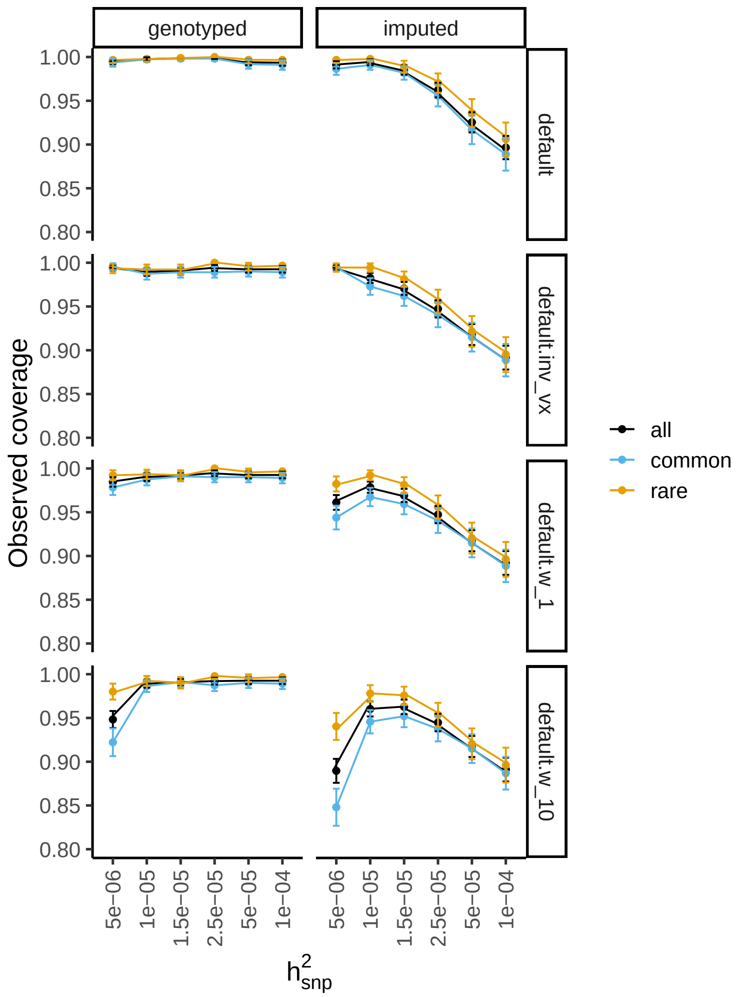

Figure S17: **The observed coverage of 95% credible sets when the focal variant is genotyped or imputed with different choices of priors.** At each $h^2_{\text{SNP}}$ (x-axis), the observed coverage (y-axis) stratified by MAF bins (common: MAF $>$ 0.01; rare: MAF $<$ 0.01) is shown. The two columns of panels show results based on genotyped and imputed values for the focal variant. For each row of the panels, the results are based on a choice of prior: 1) "default": $W = 0.1$ for all variants within the locus; 2) "default.inv_vx": $W = \frac{\text{Var}(X_\star)}{\text{Var}(X)}$ with $X_\star$ being the genotype variance of the focal variant; 3) "default.w_1": $W = 1$ for all variants within the locus; "default.w_10": $W = 10$ for all variants within the locus. Only the results based on the default method as used (i.e. no "reweighting" results).

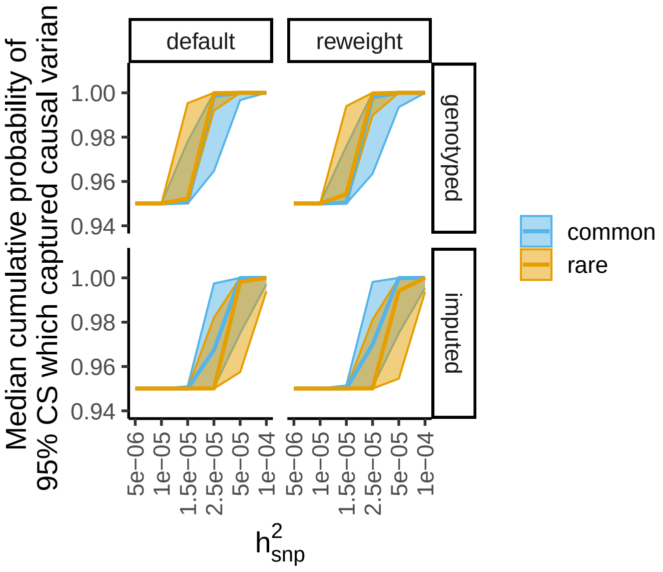

**(A)**

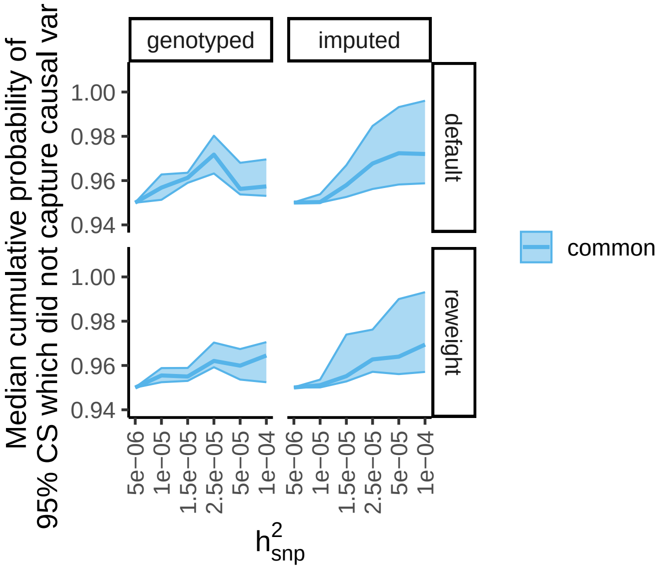

**(B)**

Figure S18: **The actual cumulative posterior probability of the 95% credible set.**. For each 95% credible set in the simulation, we calculated the actual cumulative posterior probability by summing over the PIPs across all variants with the 95% CS. The median of the actual cumulative posterior probability (thick line) under each $h^2_{\text{SNP}}$, fine-mapping method, and MAF bin (common: MAF $> 0.01$; rare: MAF $< 0.01$) are shown for credible sets capturing causal variants in panel **(A)** and credible sets not capturing causal variants in panel **(B)** in which the results on rare variants are not shown since the data is too sparse. The wide region in lighter color indicate the range between 25% to 75% quantiles of the actual cumulative posterior probability within each strata.

### 2 Supplementary Notes

#### 2.1 Derivation of the relation between $\mathbf{E}\chi^2_{\mathbf{imputed}}$ and $\mathbf{E}\chi^2_{\mathbf{genotyped}}$

Consider the widely used GWAS model $y = X\beta + \epsilon,\ \epsilon \sim N(0, \sigma^2)$ where $y$ is the phenotype and $X$ is the genotype for a variant. Consider the imputed dosage, i.e. genotype with error, being $\tilde{X} = X + e,\ e \sim N(0, \tau^2)$ (so that $\text{Cov}(X, \tilde{X}) = \text{Var}(X)$) and we define $r$ as $\text{Cor}(X, \tilde{X})$. If we assume $e \perp\!\!\!\perp X$, we have

$$
\begin{aligned}
r &:= \text{Cor}(X, \tilde{X}) && (1)\\
&= \frac{\text{Cov}(X, \tilde{X})}{\sqrt{\text{Var}(X)\text{Var}(\tilde{X})}} && (2)\\
&= \sqrt{\frac{\text{Var}(X)}{\text{Var}(\tilde{X})}},\ \text{as } \text{Cov}(X, \tilde{X}) = \text{Var}(X) && (3)
\end{aligned}
$$

The estimated effect size and $\chi^2$ statistic are

$$
\begin{aligned}
\hat{\beta} &= \frac{\widehat{\text{Cov}}(y, X)}{\widehat{\text{Var}}(X)} && (4)\\
\text{SE}(\hat{\beta}) &\approx \sqrt{\frac{\widehat{\text{Var}}(y)}{N \cdot \widehat{\text{Var}}X}}\ \text{, as per-SNP heritability is small} && (5)\\
\chi^2 &= (\frac{\hat{\beta}}{\text{SE}(\hat{\beta})})^2 && (6)\\
&\approx N \cdot \frac{\widehat{\text{Cov}}(y, X)^2}{\widehat{\text{Var}}(X)\widehat{\text{Var}}(y)} && (7)\\
&= N \cdot \widehat{\text{Cor}}(y, X)^2 && (8)\\
\text{E}\chi^2 &\approx N \cdot \text{Cor}(y, X)^2 + 1 && (9)\\
&\approx N \cdot \text{Cor}(y, X)^2 && (10)
\end{aligned}
$$

So, with genotyped values $X$ and imputed values $\tilde{X}$, we have

$$
\frac{\text{E}\chi^2_{\text{imputed}}}{\text{E}\chi^2_{\text{genotyped}}} \approx (\frac{\text{Cor}(y, \tilde{X})}{\text{Cor}(y, X)})^2 \qquad (11)
$$

$$= \underbrace{(\frac{\mathrm{Cov}(y, X+e)}{\mathrm{Cov}(y, X)})^2}_{=1 \text{ assuming } y \perp\!\!\!\perp e} \cdot \underbrace{\frac{\mathrm{Var}(X)}{\mathrm{Var}(\tilde{X})}}_{=r^2 \text{ based on Equation 3}} \quad (12)$$

$$= r^2 \quad (13)$$

#### 2.2 Implementing the "reweighting" approach in practice

Here we state the implementation details of the "reweighting" approach in practice where the input data is estimated effects $\hat{\beta}$, standard errors $\mathrm{SE}(\hat{\beta})$, LD matrix (correlation between variants) $\Sigma$, and imputation errors ($r^2$).

For numerical stability, we add an offset to the LD matrix, i.e. $\widetilde{\Sigma} = w \cdot \Sigma + (1-w) \cdot I$. Besides, any $r^2 > c$ are set to the maximum value 1 which means that these are treated as genotyped variants. We used $w = 0.1$ and $c = 0.999$ respectively in this work. Next, we calculate $\widetilde{\Sigma}^\dagger$ by calling `MASS::ginv` in `R` with `tol = 1e-5`. By slicing $\widetilde{\Sigma}^\dagger$, $S_{GI}$ and $S_{II}$ are obtained. $D_I^{-1}$ is calculated from taking all variants with $r^2 < 1$ as $D_I = \mathrm{diag}(\frac{1}{1-r^2})$. $(S_{II} + D_I^{-1})^\dagger$ is obtained by calling `MASS::ginv` with `tol = 1e-5`. The weight for the "reweighting" approach, $W_I$ (an $M \times M_I$ matrix where $M$ is the total number of variants and $M_I$ is the number of variants with $r^2 < 1$), is $\begin{bmatrix} -S_{GI} \\ D_I^{-1} \end{bmatrix} (S_{II} + D_I^{-1})^\dagger$.

Based on $W_I$, we update the z-scores for the imputed variants. For an imputed variant $j$ whose weights are $\{W_I\}_{\cdot j}$, $\tilde{z} = \frac{1}{\sigma_g} \sum_i \{W_I\}_{ij} z_i$ where $\sigma_g = \sqrt{\{W_I\}_j' \Sigma \{W_I\}_j}$. For genotyped variants, $\tilde{z} = z$. In the fine-mapping analysis, we use $\tilde{z}$ and the original estimated effect sizes as input. If the LD matrix is required, we supply a standardized version of $W' \Sigma W$ where the rows and columns are standardized so that it resembles a correlation matrix, i.e. diagonal terms are all one.

#### 2.3 Analyzing associations from UK Biobank

We downloaded ExWAS results [1] from GWAS catalog with accession numbers GCST90078986 and GCST90079277 for reported traits "Standing height (50)" and "Red blood cell erythrocyte distribution width (30070)". We downloaded GWAS summary statistics from panUKBB [2] with URLs `https://pan-ukb-us-east-1.s3.amazonaws.com/sumstats_flat_files/continuous-50-both_sexes-irnt.tsv.bgz` and `https://pan-ukb-us-east-1.s3.amazonaws.com/sumstats_flat_files/continuous-30070-both_sexes-irnt.tsv.bgz` for phenotype descriptions "Standing height" and

"Red blood cell (erythrocyte) distribution width". For panUKBB data, the results from European ancestry were used. The sample sizes for standing heights are $n_{\text{panUKBB}} = 419,596$ and $n_{\text{ExWAS}} = 430,070$. The sample sizes for red blood cell erythrocyte distribution width are $n_{\text{panUKBB}} = 408,005$ and $n_{\text{ExWAS}} = 419,192$. We obtained imputation quality measures from the variant manifest from panUKBB via URL `https://pan-ukb-us-east-1.s3.amazonaws.com/sumstats_release/full_variant_qc_metrics.txt.bgz` in which the $r^2$ was extracted from column "info". For SuSiE run, we leveraged 23andMe imputed dosages to construct the LD panel in which 50,000 customers with European ancestry were used.

To quantify the effect of imputation errors on the marginal statistics, for each of the genome-wide significant locus implicated in the panUKBB GWAS, we kept all the variants that are also present in the ExWAS with $p_{\text{ExWAS}} \geq 0.05$ and then we quantified the effect of imputation error on the association test $\chi^2$ by the slope of $\chi^2_{\text{panUKBB}} \sim \chi^2_{\text{ExWAS}}$. The slopes were obtained via the robust linear regression for model $\chi^2_{\text{panUKBB}} \sim -1 + \chi^2_{\text{ExWAS}}$ for variants with $r^2 < 0.9$ and $r^2 > 0.999$ respectively using `MASS::rlm` in `R`. The resulting coefficient of $\chi^2_{\text{ExWAS}}$, i.e. slope, represented the difference between $\chi^2_{\text{panUKBB}}$ and $\chi^2_{\text{ExWAS}}$ in the presence of various level of imputation errors.

To evaluate the quality of the 95% credible set from panUKBB standing height GWAS, we leveraged the secondary signals (via COJO [4]) of height in European ancestry from GIANT GWAS [5]. The COJO results used are from the Supplementary Table 4 of the GIANT GWAS paper [5]. We considered a credible set to be "supported" if there exists a COJO variant within $n$ bp whose maximum $R^2$ to any variants in the credible set is bigger than a cutoff $x$. We considered two cutoffs representing different stringentness: 1) $n = 100000$ and $x = 0.8$ which we referred as "supported"; 2) $n = 10000$ and $x = 0.95$ which we referred as "stringently supported". The LD panel used for getting $R^2$ was the same as the one used for SuSiE run.

##### 2.4 Analyzing the secondary signals in the F5 locus

To obtain the secondary signals from the F5 gene locus of blood clots GWAS (145,619 cases and 3,431,131 controls with dosages imputed using a panel comprising HRC, in-house WGS samples, and UK Biobank 200K WES), we performed conditional analysis (forward selection) in the 8 Mb window of the GWAS hit. At p-value < 1e-6, 11 conditional hits were identified. To isolate these secondary signals, we further performed Conditional Leave-Each-Out (CLEO) analysis which resulted in per-variant associations (CLEO statistics) after conditioning on all other conditional hits. For each secondary signal, the corresponding CLEO statistics should, in principle, capture one causal variant by conditioning on all other potential signals. Besides the primary signal, we identified three additional signals within CLEO p-value < 1e-10 laying within F5 gene body. For each of the three signals, we performed fine-mapping analysis using the CLEO statistics within 100 kb around the CLEO top variant. For the "reweighting" approach, the LD information was obtained using a random subsets of the cohort for GWAS.

#### 2.5 Summarizing the ability of capturing causal variants when imputation error appears

To summarize the effect of imputation error on fine-mapping, we consider how PIPs depend on imputation error and LD when the marginal GWAS signals take the expected $\chi^2$ and all variants have the same minor allele frequency.

For a locus with a single causal variant, let $\mathrm{E}\chi^2_\star$ denote the expected GWAS $\chi^2$ of the causal variant when genotypes are observed. Due to imputation error, the expected GWAS $\chi^2$ of the causal variants with imputed values is reduced to $r^2 \cdot \mathrm{E}\chi^2_\star$. For any other variants within the locus, their expected GWAS $\chi^2$ are $R_j^2 \cdot \mathrm{E}\chi^2_\star$ for variant $j$ where $R_j^2$ is the squared correlation between variant $j$ (observed dosages) and the causal variant (actual genotypes). The approximate Bayes factor of each variant [3] is given as follow

$$\begin{aligned}
\mathrm{PIP}_j &\propto \mathrm{ABF}_j && (14)\\
&\propto \exp(\mathrm{E}\chi^2_j) && (15)\\
&= \exp(R_j^2 \mathrm{E}\chi^2_\star) && (16)\\
&= \exp((R_j^2 - r^2)\mathrm{E}\chi^2_\star) && (17)\\
&= \exp(\Delta R_j^2 \cdot \mathrm{E}\chi^2_\star) && (18)\\
&= \underbrace{p}_{e^{\mathrm{E}\chi^2_\star}}{}^{\Delta R_j^2} && (19)
\end{aligned}$$

So,

$$\mathrm{PIP}_j \propto p^{\Delta R_j^2} \qquad (20)$$

where $\Delta R_j^2 := R_j^2 - r^2$ representing the LD to the causal variant relative to the imputation quality of the causal variant and $p = e^{\mathrm{E}\chi_\star^2}$ which is the expected GWAS $\chi^2$ of the causal variant when genotypes are observed.

Based on this result, PIPs under the expected GWAS $\chi^2$ (close to the observed PIP when GWAS power is high) depend on the LD environment of the causal variant relative to the imputation quality. Besides, the PIP is normalized value within the locus where a locus with a stronger GWAS signal ($p$ is bigger) has a sharper PIP peak comparing to the same locus ($\Delta R_j^2$ is the same) with a weaker GWAS signal ($p$ is smaller). Furthermore, the shape of the PIP peak also depends on the distribution of $R_j^2$ which relates to haplotype length.

To make a systematic illustration of the ability of 95% credible set on pinpointing the causal variants with different LD environments, haplotype lengths, and imputation qualities, we considered a simulated toy locus. This locus contains 2000 variants in which $K$ variants are in high LD ($R^2_{\text{high}}$ varying from 0.1 to 0.99) with the causal variant and $2000 - K$ variants are in low LD ($R^2_{\text{low}} = 0.1$) with the causal variant. $K$ varied from 5 to 500 to represent a range of haplotype lengths. The causal variant has imputation quality $r^2$ varying from 0.3 to 1. We consider a wide range of GWAS power (at the causal variant, i.e. $\mathrm{E}\chi_\star^2$) from p-value = 1e-6 to 1e-100 and use Equation 20 to calculate PIPs.

In Figure S16, we show the minimum imputation $r^2$ required to have the 95% credible set capture the causal variant. This result suggests that a better powered GWAS, a locus with higher LD, or a locus with longer haplotypes all require a more well-imputed causal variant to be able to capture the causal variant in the 95% credible set. These observations nicely summarize what we observed in our simulation study and real data case study and provide additional insights on dissecting the effect of imputation errors in various scenarios.
